# Experimental colitis drives enteric alpha-synuclein accumulation and Parkinson-like brain pathology

**DOI:** 10.1101/505164

**Authors:** Stefan Grathwohl, Emmanuel Quansah, Nazia Maroof, Jennifer A. Steiner, Liz Spycher, Fethallah Benmansour, Gonzalo Duran-Pacheco, Juliane Siebourg-Polster, Krisztina Oroszlan-Szovik, Helga Remy, Markus Haenggi, Marc Stawiski, Matthias Sehlhausen, Pierre Maliver, Andreas Wolfert, Thomas Emrich, Zachary Madaj, Martha L. Escobar Galvis, Christoph Mueller, Annika Herrmann, Patrik Brundin, Markus Britschgi

**Affiliations:** Roche Pharma Research and Early Development, Neuroscience Discovery, Roche Innovation Center Basel, F. Hoffmann-La Roche Ltd, Grenzacherstrasse 124, Basel, Switzerland; Center for Neurodegenerative Science, Van Andel Research Institute, 333 Bostwick Ave. NE, Grand Rapids, MI, USA; Roche Pharma Research and Early Development, pREDi, Roche Innovation Center Basel, F. Hoffmann-La Roche Ltd, Grenzacherstrasse 124, Basel, Switzerland; Roche Pharma Research and Early Development, Pharmaceutical Sciences, Roche Innovation Center Basel, F. Hoffmann-La Roche Ltd, Grenzacherstrasse 124, Basel, Switzerland; Roche Pharma Research and Early Development, Pharmaceutical Sciences, Roche Innovation Center Munich, Roche Diagnostics GmbH, Nonnenwald 2, Penzberg, Germany; Institute of Pathology, University of Bern, Murtenstrasse 31, Bern, Switzerland

## Abstract

Intraneuronal α-synuclein accumulation is key in Parkinson’s disease (PD) pathogenesis. The pathogenic process is suggested to begin in the enteric nervous system and propagate into the brain already decades before diagnosis of PD. In some patients, colitis might play a critical role in this process. Here we demonstrate that patients with inflammatory bowel disease exhibit α-synuclein accumulation in the colon and that experimental colitis triggers α-synuclein accumulation in certain enteric nerves of mice. The type and degree of experimental inflammation modulates the extent of colonic α-synuclein accumulation and macrophage-related signaling limits this process. Remarkably, experimental colitis at three months of age exacerbates the accumulation of aggregated phospho-Serine 129 α-synuclein in the midbrain (including the substantia nigra), in 21- but not 9-month-old α-synuclein transgenic mice. This is accompanied by loss of tyrosine hydroxylase-immunoreactive nigral neurons. Our data suggest that intestinal inflammation might play a critical role in the initiation and progression of PD.

---

Parkinson’s disease (PD) is a progressively debilitating neurodegenerative disease affecting 1% of the population above 60 years ^1^. Typical symptoms are motor impairments including muscle rigidity, tremor, and bradykinesia. Neuropathologically, PD is hallmarked by loss of dopaminergic neurons in the substantia nigra (SN), a concomitant reduction of striatal dopaminergic signaling ^2^, and the presence of intraneuronal inclusions called Lewy bodies and neurites ^3^. Lewy pathology is enriched in α-synuclein (αSyn), a presynaptic protein that tends to aggregate and become phosphorylated under pathological conditions ^2^. Rare point mutations in αSyn and gene multiplications also cause familial forms of PD and related neurological conditions, and certain single nucleotide polymorphisms close to the αSyn gene (*SNCA*) locus are associated with increased risk for sporadic PD ^4^. These findings make αSyn a focal point of biomarker and drug development programs for PD.

Several years before the first appearance of motor symptoms, many patients exhibit a variety of non-motor symptoms including constipation, sleep disorder, depression, and hyposmia ^5–7^. Indeed, co-occurrence of some of these non-motor symptoms is coupled to elevated PD risk ^8–11^. Constipation is an important non-motor feature of prodromal PD, with 28-61% of patients having exhibited gastrointestinal dysfunction for several years during the prodrome ^7,10,12^. Notably, αSyn-immunoreactive inclusions have been found in neurons of the submucosal plexus in people with PD ^3,13^. Taken together, this converging evidence suggests an early involvement of the enteric nervous system (ENS) in the pathogenesis of PD. Already over a decade ago, Braak and colleagues hypothesized that αSyn-immunoreactive inclusions first appear in the ENS and then gradually engage the brainstem, including the dorsal motor nucleus of the vagus nerve and midbrain areas ^3,13^. Several studies in preclinical models have demonstrated that αSyn pathology in the gut is associated with the development of αSyn pathology in the brain ^14–18^. It will be critical to determine factors that regulate αSyn accumulation in the ENS and to understand whether the process underlying αSyn accumulation in the gut can also lead to αSyn pathology in the brain.

Inflammation can potentially trigger αSyn pathology in the ENS of the gut and in the brain. A recent finding in children with gastrointestinal inflammation suggests an immune regulatory function of αSyn ^19^. Immune pathways are indeed activated in the brain and colon of PD cases ^20,21^. Also, several genes associated with an increased PD risk have an immune system-related function ^22^, and it was recently proposed that PD heritability is not simply due to variation in brain-specific genes, but that several cell types in different tissues might be involved ^23^. Adding further genetic evidence supporting that inflammation is involved in PD pathogenesis, a genome-wide association study identified common genetic pathways linking PD and autoimmune disorders ^24^. Most prominently, LRRK2, a major genetic risk factor for PD also confers increased risk for developing inflammatory bowel disease (IBD) ^25^ and is known to modulate the function of monocytes, macrophages and other immune cells ^26,27^. Intriguingly, IBD is associated with an increased risk for developing PD and specifically blocking the TNF pathway reduces this risk ^28–31^. This suggests that the intestinal immune environment plays a role in triggering PD or facilitating the molecular events involved in the earliest phases of the disease process ^32^.

Here we tested the hypothesis that intestinal inflammation (e.g. colitis) triggers accumulation of αSyn in the ENS and the subsequent development of αSyn pathology in the brain. We discovered that patients with IBD exhibited increased αSyn accumulation in the submucosa of the colon. Experimental forms of colitis in wild type and αSyn transgenic mice demonstrated that the type and degree of inflammation regulates the amount of αSyn accumulation in the colon. Macrophage-related signaling limited the extent of αSyn immunoreactivity. When αSyn transgenic mice were exposed to experimental colitis at 3 months of age and then were aged normally up to 9 or 21 months, the accumulation of aggregated αSyn in midbrain, including the SN, was much exacerbated in the 21-months age group, but not in the 9-months age group. These 21-month old mice also exhibited loss of nigral tyrosine hydroxylase-immunoreactive neurons. Together, our data support a critical role for intestinal inflammation in the initiation and progression of PD.

## Results

### IBD patients show αSyn accumulation in the ENS and local macrophages

Recent epidemiological data links inflammatory bowel disease (IBD) to an increased PD risk ^28–30^. In order to explore if IBD is associated with enteric αSyn accumulation we performed immunohistochemistry for αSyn in cryo-sections from colonic biopsies of patients with ulcerative colitis (UC, n = 11, mean age 31 years), Crohn’s disease (CD, n = 11; mean age 35 years), and from healthy subjects (HS, n = 8; mean age 51 years). We observed in eight UC cases various degrees of αSyn accumulation, mostly in structures with the morphology of neurites (Figure 1). Interestingly, the eight UC cases, and four patients with CD (images not shown) also showed marked intracellular αSyn staining in many infiltrating monocytic cells. In contrast, only one HS showed a few cells immunoreactive for αSyn (images not shown). This finding in human tissue suggests a potential role of local inflammation in the development of enteric αSyn accumulation.

**Figure 1.**
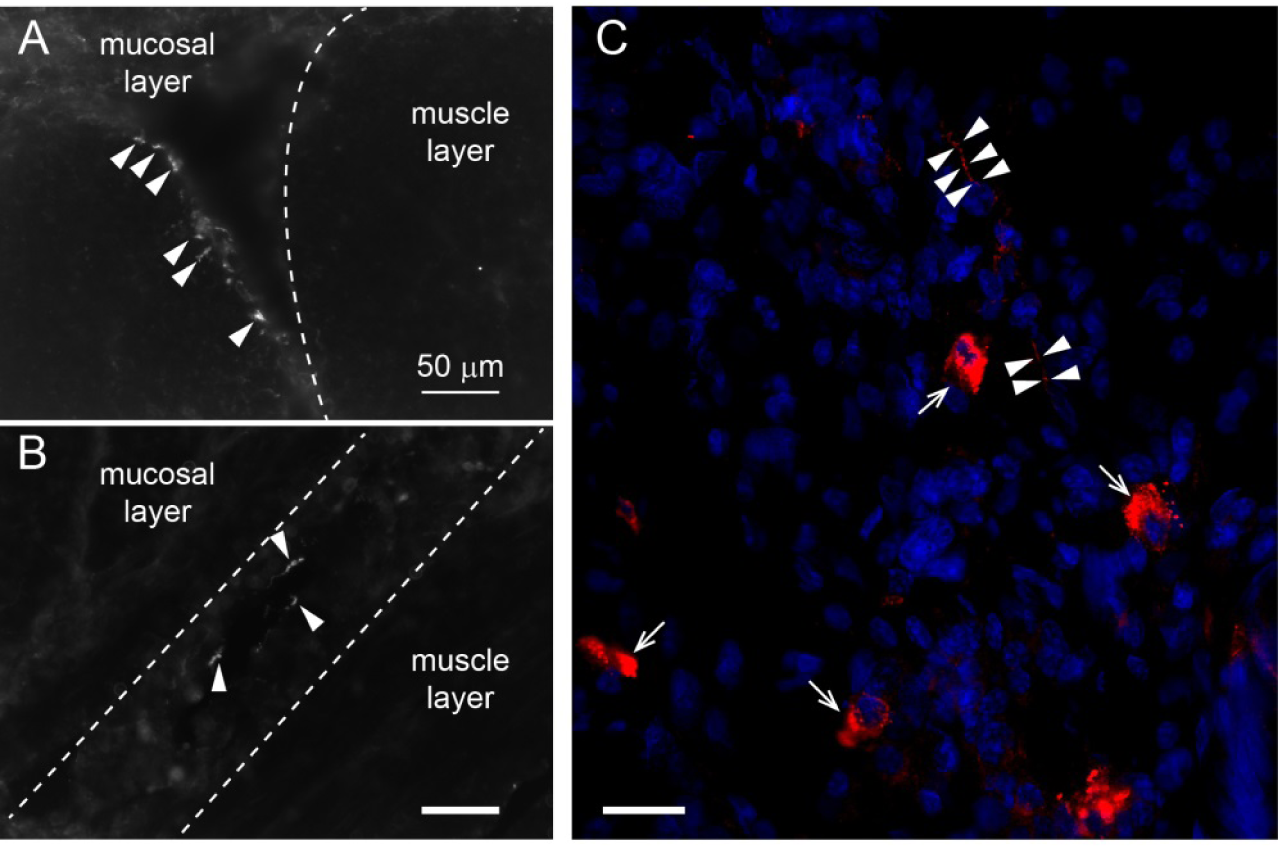
Alpha-Synuclein inclusions in the enteric nervous system and in macrophages of patients with inflammatory bowel disease. **(A, B)** Immunofluorescence images of αSyn inclusions (Syn1 antibody) in the submucosal region of 10 μm cryo-sections from colons of patients with colitis ulcerosa. Arrow heads point to neuritic features indicating presence of inclusions in enteric nerves. Scale bar 50 μm. **(C)** Close-up of a colonic region with active leukocyte infiltration in a patient with ulcerative colitis. Immunoreactivity for αSyn (red) was observed in neuritic features (arrow heads) as well as in individual leukocytes (arrows; identification of leukocytes based on cellular morphological features and localization). Nuclei are shown in blue (DAPI). Scale bar 20 μm.

### Experimental IBD exacerbates αSyn load in submucosal plexus of αSyn transgenic and wildtype mice

During the process of further characterizing a (Thy1)-h[A30P]αSyn transgenic mouse line ^33^ we detected human αSyn accumulation in all innervated organs that we analyzed (**Supplemental Figure 1**). This included the myenteric and submucosal plexuses of the ENS, where human αSyn co-localized with peripherin, a specific marker for peripheral nerves (Figure 2A). We observed an age-dependent increase of baseline human αSyn inclusions (irregularly sized and shaped inclusion bodies detected by human αSyn specific monoclonal antibody clone 211) in both plexuses between the ages of three and twelve months (Figure 2B). We wanted to test whether IBD-related inflammation in the colon exacerbates this local accumulation of αSyn acutely (e.g. within a few days or weeks) and how the age of the αSyn transgenic mice influenced the outcome. Administration of dextran sulfate sodium (DSS) in the drinking water in acute or chronic paradigms are well-established mouse colitis models of IBD, exhibiting infiltration of leukocytes into the submucosa with various degrees of destruction of the colonic mucosa and submucosa ^34^. Due to awareness of the variability of the DSS model in different genetic backgrounds of mice, we first tested DSS administration at different concentrations and durations in the (Thy1)-h[A30P]αSyn transgenic mice (Figure 2C), and observed leukocyte infiltration in a dose-dependent manner and which was similar at the age of 3 and 6 months (Figure 2D **and** 3A). In the acute paradigm with mice at the age of 3 months, 2.5%, but not 1%, DSS triggered intracellular accumulation of αSyn in nerves of the submucosal plexus (Figure 3A, B). In the chronic DSS paradigm, which was done with mice at the age of 6 months, we observed a dose-dependent increase of αSyn load in the submucosal plexus, but at a smaller magnitude than in the younger mouse cohort (Figure 3A). Wildtype mice also express endogenous αSyn in innervated organs, but at much lower levels compared with the overexpressed human αSyn protein in the heterozygous (Thy1)-h[A30P]αSyn transgenic mice (**Supplemental Figure 1**). To confirm that the finding in (Thy1)-h[A30P]αSyn transgenic mice was independent of transgenic expression of human αSyn, we applied acute and chronic (consistent dose) DSS paradigms also in wildtype mice. In both treatment paradigms, we observed an elevated number of inclusion bodies of endogenous murine αSyn in the submucosal plexus (detected by rodent cross-reactive αSyn-specific monoclonal antibody Syn1/clone 42, Figure 3C, D). A separate experiment also confirmed that the observed effects of DSS could not be attributed to increased gene expression of murine or the transgenic human αSyn **(Supplemental Figure 2)**. Together, these results confirmed the validity of this experimental IBD paradigm to test the effect of inflammation on αSyn accumulation in the ENS in wild type and (Thy1)-h[A30P]αSyn transgenic mice. Because 3-month old (Thy1)-h[A30P]αSyn transgenic mice provided more optimal conditions for visualization and quantification of αSyn inclusions in the ENS, for the remainder of the study we focused on using this transgenic mouse model.

**Figure 2.**
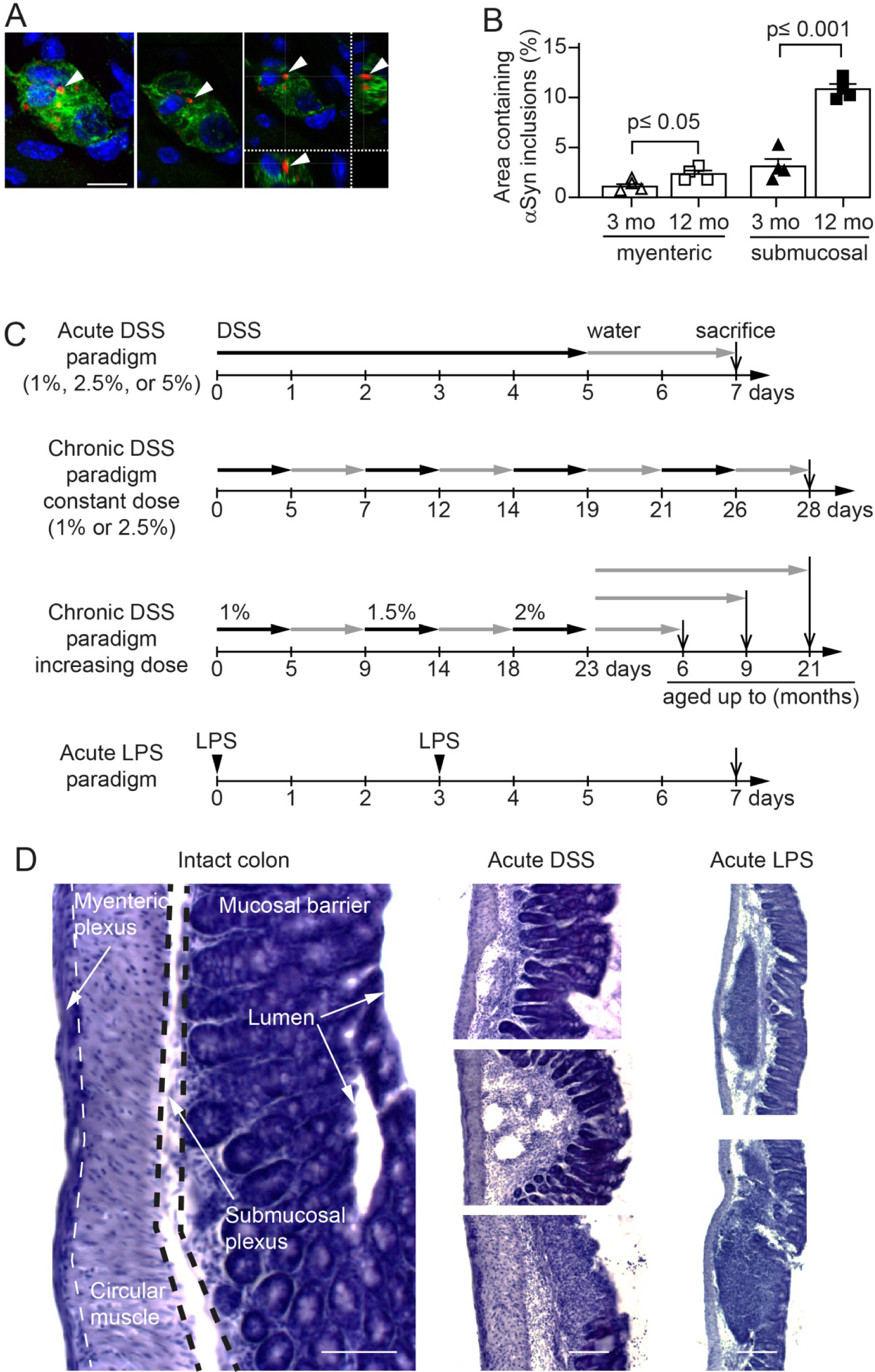
Age dependent increase of intracellular αSyn accumulation in enteric nervous system of heterozygous (Thy1)-h[A30P]αSyn transgenic mice and setup of the experimental colitis paradigms. **(A)** Confocal microscopy imaging of the inclusions of human αSyn (red, antibody clone 211; human αSyn specific) within the ganglia of the submucosal plexus (green, peripherin; blue, DAPI/nuclei) of heterozygous (Thy1)-h[A30P]αSyn transgenic mice. Arrow head points to one of the typical irregularly sized and shaped αSyn inclusion bodies visualized in 2D z-stacks of rotated confocal images. Scale bar, 100 μm. **(B)** Stereological quantification of normally occurring human αSyn inclusions in the myenteric and submucosal plexuses of 3 and 12 months old heterozygous (Thy1)-h[A30P]αSyn transgenic mice (n = 4 per group; mean and S.E.M. are shown; Student t-test between the two age groups in each region). **(C)** Setup of experimental colitis paradigms employing dextran sulfate sodium (DSS, per os in drinking water). Additionally, peripheral inflammation was induced by bacterial lipopolysaccharide (LPS, intraperitoneal injection). After some chronic DSS paradigms mice were aged on normal water up to 6, 9 or 21 months. Mice aged up to 9 or 21 months of age were analyzed for brain pathology **(D)** Hematoxylin staining of 35 μm thick colon sections of 3 months old heterozygous (Thy1)-h[A30P]αSyn transgenic mice. Organizational layers of the intact colon (left panel). Representative images of various severity degrees of DSS-driven colitis from weak leukocyte infiltration (top panel of acute DSS) to mucosal ulceration (lowest panel of acute DSS). Note the different appearance of enteric inflammation in acute LPS-driven peripheral inflammation compared with DSS; e.g., confined immune cell clustering and lymphoid hyperplasia; intact mucosal layer. Scale bar 50 μm (intact colon), 100 μm (acute DSS), and 200 μm (LPS).

**Figure 3.**
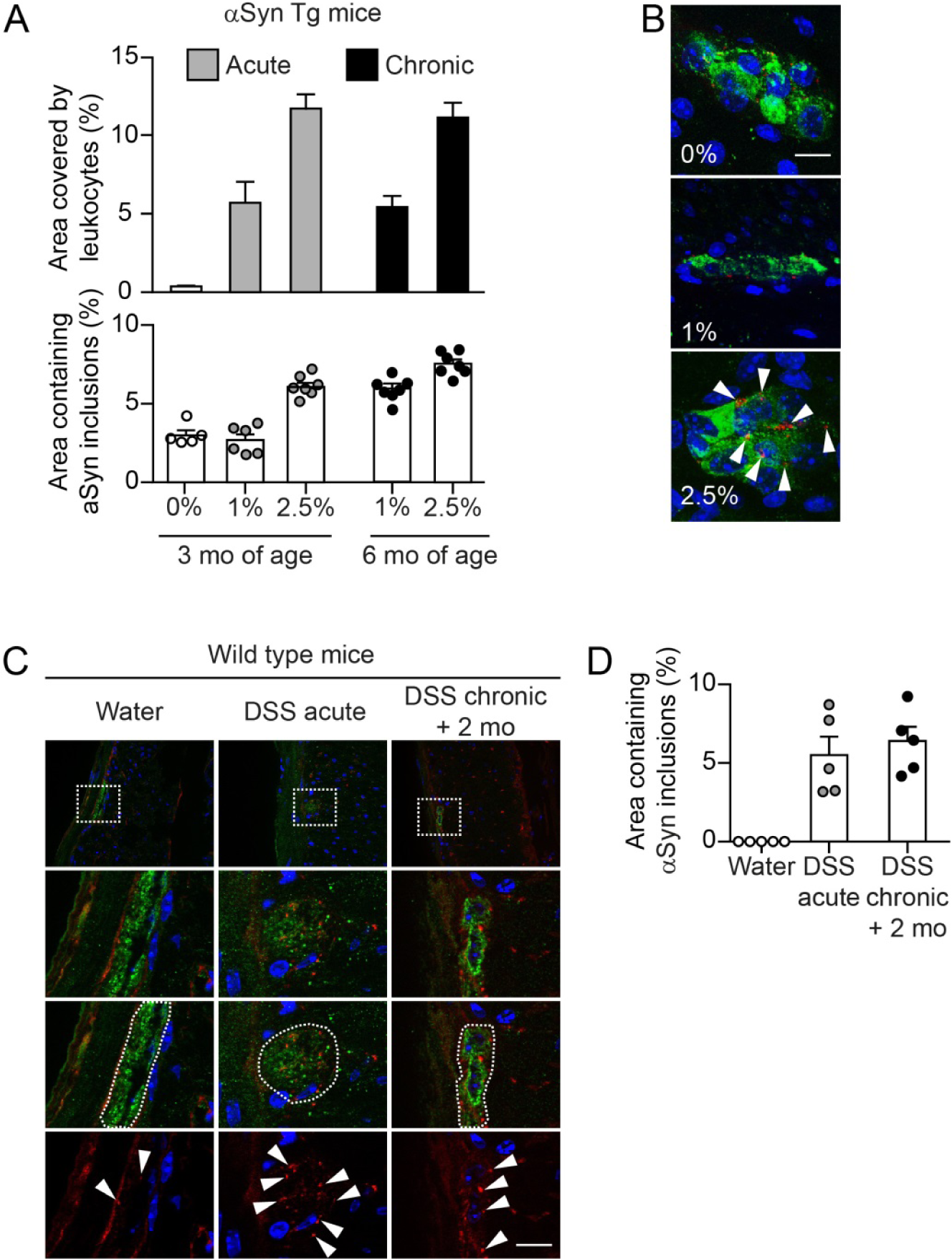
Colitis severity and duration-dependent aggravation of accumulation of αSyn inclusions in the colonic submucosal plexus of heterozygous (Thy1)-h[A30P]αSyn transgenic and wild type mice. **(A)** DSS dose-dependent increase of leukocyte infiltration in the acute and chronic paradigm. The highest acute dose (2.5%) and the two constant chronic doses led to an increase of αSyn inclusions in the submucosal plexus (stereological quantification of αSyn inclusions in the submucosal plexus of 3 and 6 months old heterozygous (Thy1)-h[A30P]αSyn transgenic mice; n = 5-7 per group; mean and s.e.m. are shown). **(B)** Representative 2D z-stacks of confocal images of increasing abundance of αSyn inclusions (red, human-αSyn specific monoclonal antibody clone 211) in a ganglion of the submucosal plexus (green, peripherin) with cellular nuclei in blue (DAPI) in the acute DSS paradigm. Arrow heads point to the typical irregularly sized and shaped αSyn inclusion bodies that accumulate in the highest DSS dose. Scale bar 200 µm. **(C)** Overview of colonic region of 3-month-old wildtype mice (top row) exposed to water or acute DSS (5%) with immunofluorescence analysis of murine αSyn load in the colon performed immediately after colitis or exposed to constant chronic DSS (2.5%) and analysis after aging on normal water for another 2 months. White dotted rectangles in the top row indicate the area that was zoomed out below illustrating in more detail the murine αSyn inclusions (red, rodent αSyn cross-reactive monoclonal antibody syn1/clone 42) in the submucosal plexus (green, peripherin). The lower three rows shows DAPI and αSyn inclusions with and without the peripherin channel. The white dotted circled area illustrates the peripherin-positive area that was analyzed for αSyn inclusion bodies (arrow heads in bottom row). Scale bar for the lower three panels 200 μm. **(D)** Stereological quantification of murine αSyn inclusions in the submucosal plexus of wildtype mice right after acute DSS colitis or after 2 months of recovery from a 4-week chronic DSS colitis (n = 5 per group). Note the regularly arranged and smoothly distributed immunoreactivity for the physiological αSyn with barely any inclusion bodies in the intact enteric nerves of the water group.

### Colitis induced by peroral DSS but not by peritoneal administration of LPS aggravates αSyn accumulation in colonic submucosal plexus of αSyn transgenic mice

In order to explore effects of different approaches to induce inflammation in or nearby the gut in (Thy1)-h[A30P]αSyn transgenic mice, we compared the outcome of acute 5% DSS in drinking water with acute 0.5 mg/kg intraperitoneal LPS administration (Figure 2C **and** 4). In order to maximize the inflammatory response, we administered both DSS and LPS at high, but still tolerable, doses. At day 7, both agents had induced variable degrees of leukocyte infiltration in the submucosa of the colon while a marked destruction of the mucosa was induced when giving only DSS (Figure 2D). As before, the DSS-exposed mice presented with increased accumulation of αSyn in the ganglia of the submucosal plexus (Figure 4A). In contrast, we detected no change in αSyn load in the myenteric plexus, consistent with lack of leukocyte infiltration in this part of the colonic wall (Figure 4B).

**Figure 4.**
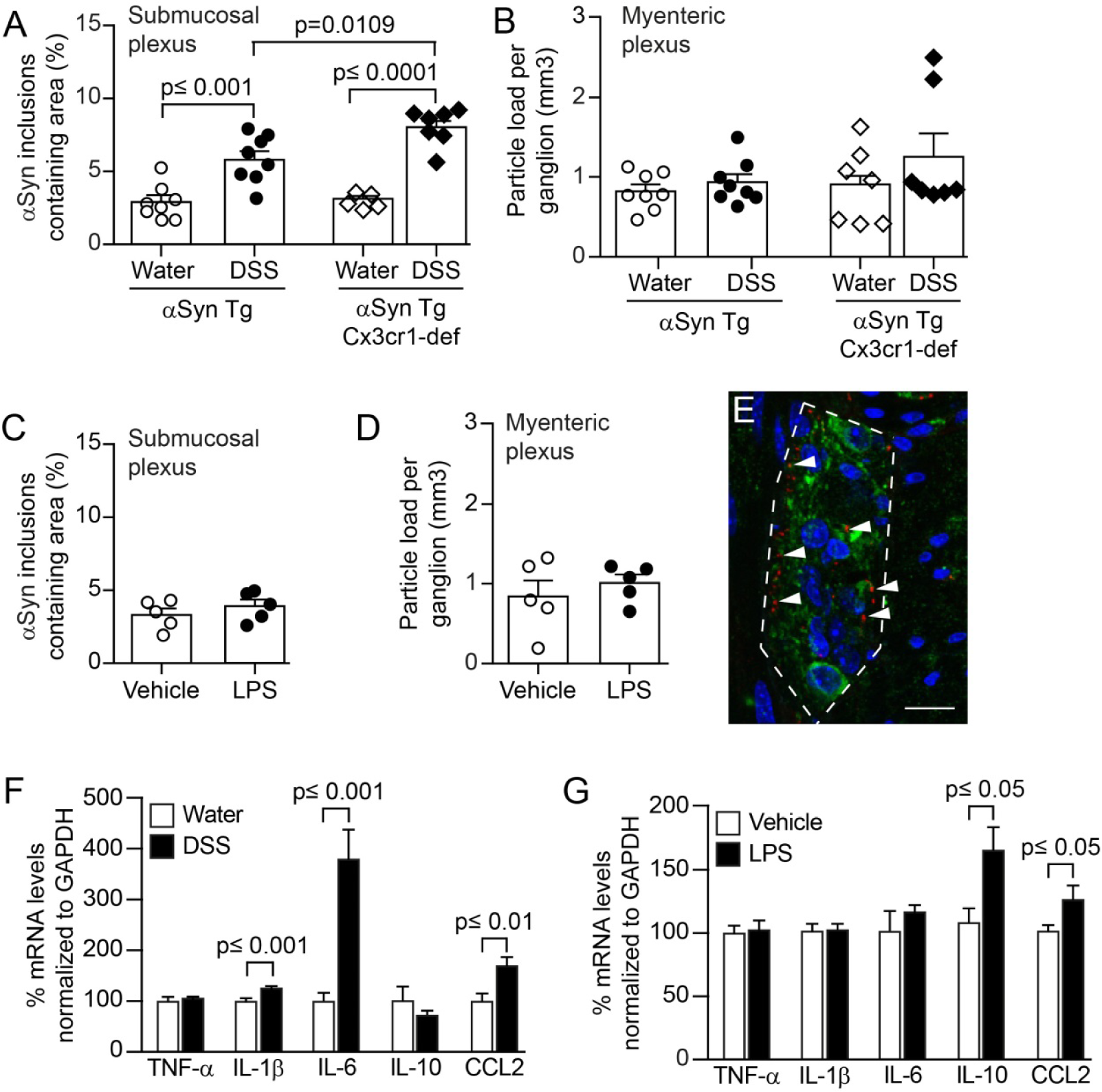
Colitis induced by peroral DSS but not peritoneal LPS enhances αSyn accumulation in the colonic submucosal plexus of heterozygous (Thy1)-h[A30P]αSyn transgenic mice and can be aggravated by lack of Cx3cr1 signaling. Mice received in an acute paradigm either peroral 5% DSS in their drinking water or intraperitoneally 0.5 mg/kg LPS. Effects of the two agents in the ENS was compared to effects induced by vehicle (see Figure 2C for timelines). Stereological quantification of αSyn inclusions in the submucosal plexus as % area **(A, C)** and in the mucosal plexus as particle load per ganglion **(B, D)** (Two-way ANOVA with Tukey post hoc test). **(E)** Representative 2D stacks of confocal images of intracellular αSyn inclusions (red, human αSyn specific monoclonal antibody clone 211; arrow heads pointing to some selected inclusions) in a ganglion of the myenteric plexus (green, peripherin) with cellular nuclei in blue (DAPI). Scale bar 50 μm. Gene expression analysis of selected cytokines in the colon of (Thy1)-h[A30P]αSyn transgenic mice that received either acutely LPS **(F)** or DSS **(G)** compared to their respective vehicle or water controls. Note the strong increase in IL-6 and the lack of elevation of IL-10 in the DSS paradigm compared to the LPS paradigm indicating a different inflammatory colonic milieu despite the abundant leukocyte infiltration in both paradigms. n = 5-8 per group; mean and s.e.m.; Student’s t-test between inflammatory agent and vehicle for individual cytokines.

Despite the high dose, LPS-induced inflammation did not increase αSyn accumulation in the colonic nervous plexuses (Figure 4C, D). Notably, LPS and DSS resulted in a differential expression of cytokines, and consistent with leukocyte recruitment, CCL2 was elevated in both (Figure 4F, G). In the LPS paradigm, mRNA for IL-10 was markedly elevated, whereas DSS strongly increased IL-6 and also IL-1β but not IL-10. Together these results indicate that only certain types of local inflammation increase the intracellular accumulation of αSyn in the colon.

### Lack of Cx3cr1 signaling during DSS colitis aggravates αSyn load in the submucosal plexus of ***αSyn transgenic mice***

In both the IBD patients and the (Thy1)-h[A30P]αSyn transgenic mice that experienced acute DSS colitis, we observed several αSyn-positive cells with a morphology consistent with them being infiltrating leucocytes (Figure 5). In the mice, these infiltrating cells were positive for the macrophage marker Iba-1 (Figure 5C-D). In order to explore further the role of monocytes/macrophages in the accumulation of αSyn in our DSS model, we added an experimental arm with (Thy1)-h[A30P]αSyn transgenic mice crossed with mice that have a deletion of Cx3cr1 induced by an insertion of GFP (Cx3cr1-GFP knock-in mice) (Figure 4A, B). The CX3CR1-CX3CL1 axis plays an important role in maintaining the function of the lamina propria macrophage population of the gastrointestinal wall and lack of this signaling pathway in experimental colitis models may either aggravate or ameliorate the induced pathology ^35–37^. In our experiment, the area covered by infiltrating leukocytes following exposure to DSS located to the mucosa and submucosa and was not significantly higher in the Cx3cr1-deficient αSyn transgenic mice than in the Cx3cr1-competent mice (**Supplemental Figure 2A**). However, a significantly higher level of αSyn accumulated in the submucosal plexus in αSyn transgenic mice lacking Cx3cr1 compared to αSyn transgenic mice expressing Cx3cr1 (p = 0.001, two-way ANOVA with Tukey HSD post-hoc analysis; Figure 4A). In the myenteric plexus, we found no significant increase in αSyn accumulation in neither the αSyn transgenic mice with normal Cx3cr1 nor the αSyn transgenic mice deficient in Cx3cr1, indicating again a prominent role for the localization of leukocyte infiltration in the process of αSyn accumulation in the submucosa (Figure 4B). Collectively, our results indicate a potential link between monocyte/macrophage signaling and αSyn accumulation in ENS in this experimental IBD model.

**Figure 5.**
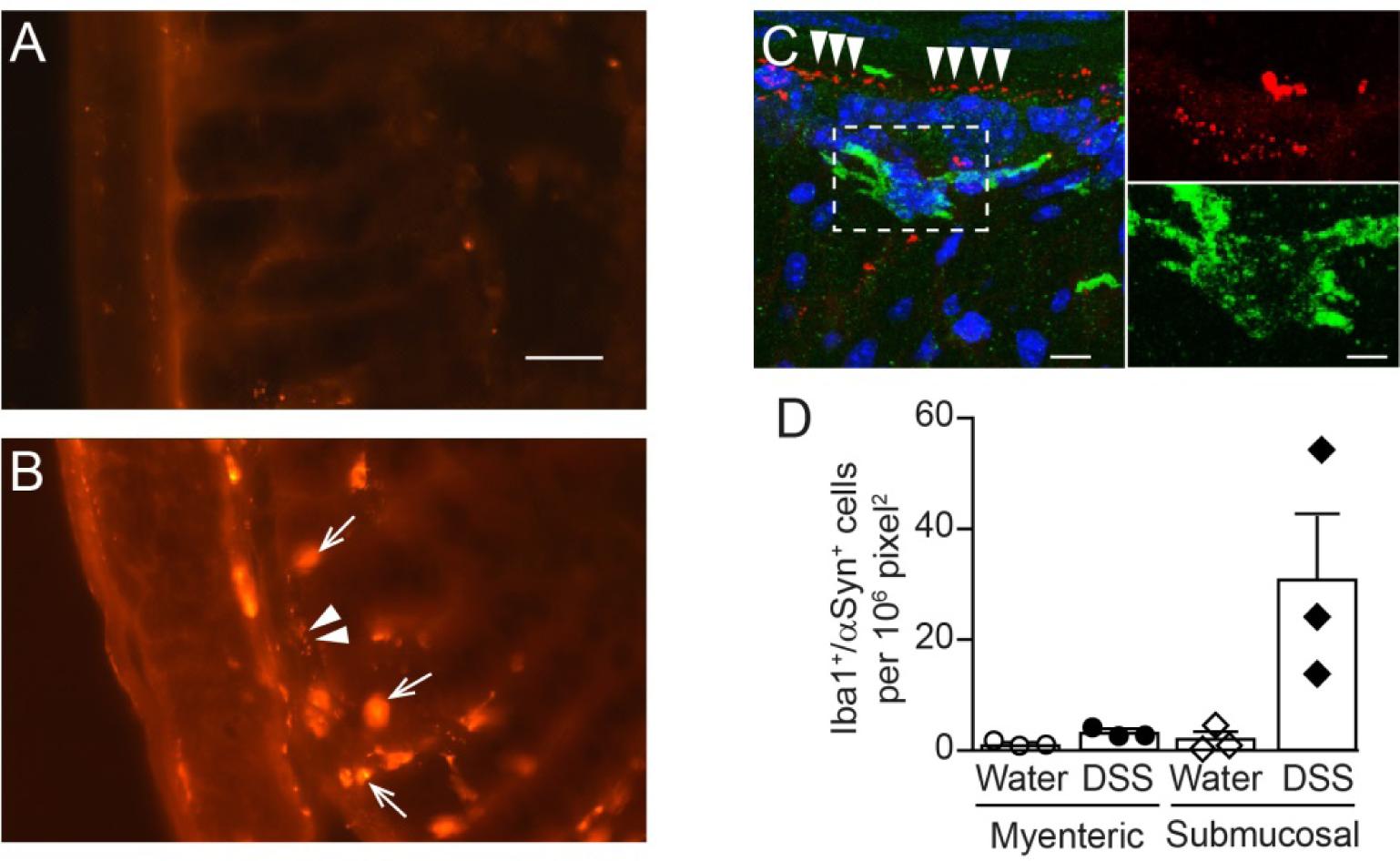
Alpha-synuclein co-localizes with ENS and macrophages upon DSS colitis in αSyn transgenic mice. **(A, B)** Immunofluorescence image of αSyn staining in colonic region of (Thy1)-h[A30P]αSyn transgenic mice on water **(A)** or after acute DSS colitis (2.5%) **(B)**. Note the small dotted structures of the typical αSyn inclusions in the submucosal plexus (arrow heads) and the large features of immunoreactivity which localize to infiltrating leukocytes (arrows; identified by their typical cellular morphology), similar to what was observed in IBD patients in Figure 1. Scale bar 100 µm. **(C)** 2D stacks and close-up of confocal images co-localizing αSyn (red) with the macrophage marker Iba-1 (green) in the colon of a (Thy1)-h[A30P]αSyn transgenic mouse after DSS colitis. Note the dotted structures of the typical αSyn inclusions in the submucosal plexus (arrow heads). Scale bar 40 μm and 13 μm for the close-up. **(D)** Quantification of numbers of Iba-1/αSyn-double positive macrophages (n = 3 per group; mean and S.E.M.)

### Systemic IL-10 ameliorates DSS-induced colitis and associated enteric αSyn accumulation in αSyn transgenic mice

As mentioned above, LPS-induced colonic leukocyte infiltration did not result in increased accumulation of αSyn in the ENS of the colon and that the main difference in cytokine expression between the DSS and LPS paradigms was increased IL-10 expression in the LPS group (Figure 4). Interleukin-10 is an important regulator of monocytes/macrophages, and genetic ablation of IL-10 signaling or blocking IL-10 with specific antibodies has been reported to enhance DSS colitis ^38,39^. To mimic the effect of higher levels of IL-10 in an acute model of DSS colitis (5% DSS, Figure 2C), we administered intravenously murine IL-10 (mIL10), which was recombinantly engineered onto two different murine IgG variants to extend the half-life of mIL-10 in circulation (mIgG1(v1)-mIL10 and mIgG1(v2)-mIL10, respectively). As described above, DSS induced a marked increase in leukocyte infiltration and αSyn accumulation, and we found this to be similar in the untreated and control IgG treated group (Figure 6A, B). By contrast, both mIgG1(v1)-mIL10 and mIgG1(v2)-mIL10 significantly reduced leukocyte infiltration in mice treated with DSS (p<0.0001, one-way ANOVA with Tukey HSD post-hoc analysis; Figure 6A, B). Regarding human αSyn in the submucosal plexus, only mIgG1(v2)-mIL10 significantly reduced the levels in DSS treated mice (p=0.02, one-way ANOVA with Tukey HSD post-hoc analysis; Figure 6B). Interestingly, the significantly reduced αSyn accumulation was associated with detectable serum exposure of mIgG1(v2)-mIL10, whereas mIgG1(v1)-mIL10 was no longer detectable at the end of the *in vivo* phase, after 7 days (Figure 6C). These results underline further an important role for the IL-10 pathway in keeping αSyn accumulation at a reduced level throughout the course of experimental IBD. Together, our results with the genetic (e.g., CX3CR1-CX3CL1 axis) and pharmacological modulation (e.g., IL-10) of DSS colitis corroborate an important role for monocyte/macrophage pathways in the development of αSyn accumulations in the ENS of the colon.

**Figure 6.**
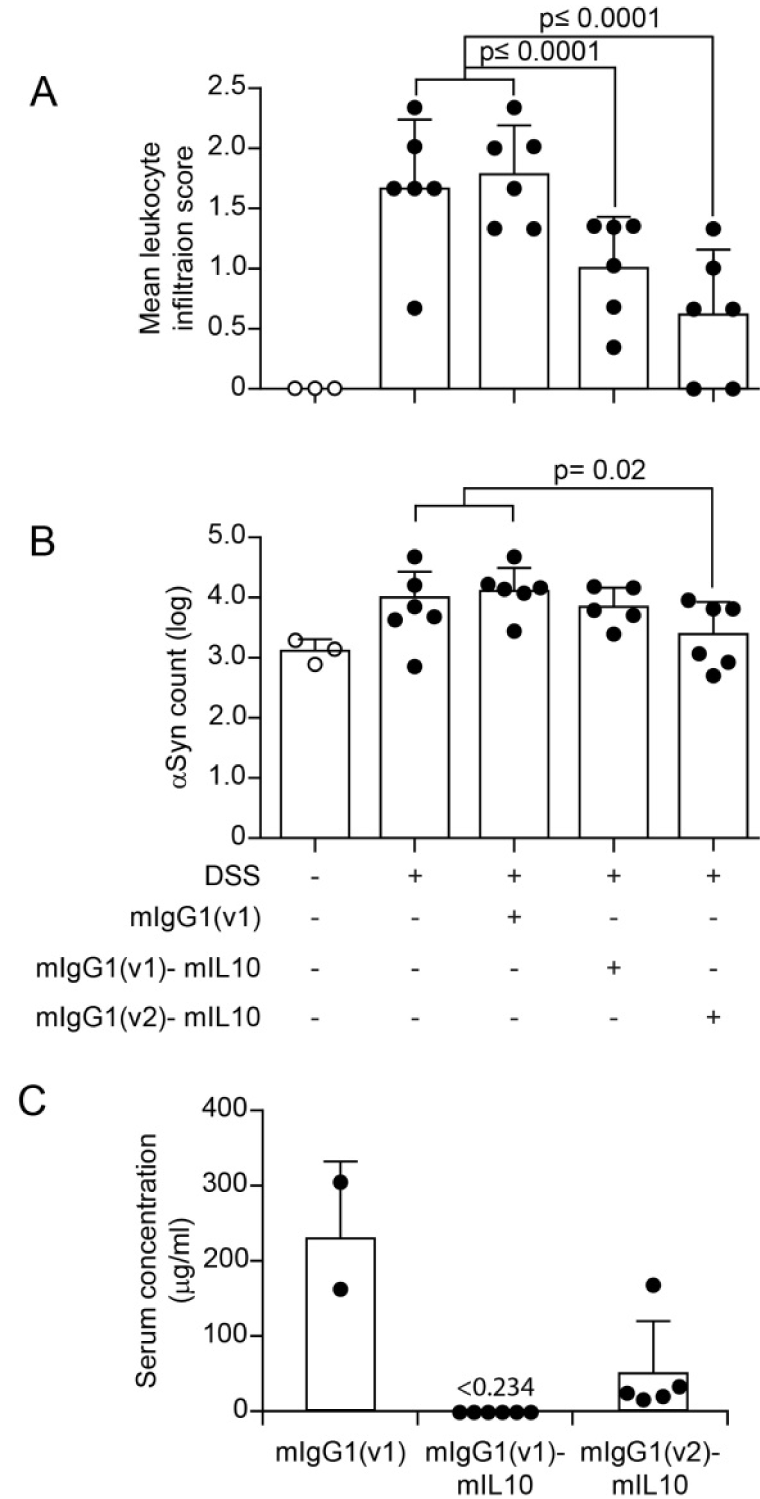
Systemic IL-10 ameliorates DSS colitis and associated local αSyn accumulation in (Thy1)-h[A30P]αSyn transgenic mice. Two different recombinantly engineered and murine IgG1-fused forms of murine IL-10 (mIgG1(v1)-mIL10 and mIgG1(v2)-mIL10) were administered (150 µg per mouse i.p.) at the beginning of the acute DSS paradigm (5%) in (Thy1)-h[A30P]αSyn transgenic mice. Vehicle and the mIgG1(v1) alone served as untreated controls. **(A)** Leukocyte infiltration was assessed by visual scoring and **(B)** inclusion features of αSyn were stereologically and semi-automatically quantified and result log scaled for statistical analysis. Both the vehicle group and the mIgG1(v1) group had similar levels of leukocyte infiltration and αSyn inclusions and were merged for the statistical analysis to compare with the IL-10 treated groups. Both forms of IL-10 ameliorated leukocyte infiltration whereas mIgG1(v2)-mIL10 also blocked the appearance of αSyn inclusions significantly (n = 3-6 per group; mean and s.e.m.; one-way ANOVA and Tukey post hoc test). **(C)** Persistent exposure mIgG1(v2)-mIL10 versus mIgG1(v1)-mIL10 (lower limit of detection is indicated at <0.234 μg/ml) as measured in serum at the end of the in vivo phase corresponds with beneficial treatment effects on αSyn readout observed above. The mIgG1(v1) was only measured in two mice.

### DSS colitis-induced submucosal αSyn accumulation at a young age persists for months and is exacerbated by lack of Cx3cr1 signaling

IBD increases PD risk ^28–30^ and our own data in colon samples from IBD patients (Figure 1) and recent evidence in Crohn’s disease ^40^ indicate that such gut inflammatory conditions are associated with αSyn accumulation in the ENS ^31^. Because longer exposure to DSS mimics more closely the chronic nature of IBD, we next elected to explore αSyn accumulation in the submucosal plexus of (Thy1)-h[A30P]αSyn transgenic mice that were subjected to DSS colitis in a 4-week chronic increasing dose paradigm. In order to allow for a full recovery from the chronic inflammation we then left the mice for two months on normal drinking water and analyzed them at the age of 6 months (Figure 2C). Because we also wanted to explore the effect of modulating monocytes/macrophages in this chronic setting, an experimental arm with (Thy1)-h[A30P]αSyn transgenic mice crossed with Cx3cr1-deficent mice was added. As expected, the area that is usually extensively covered by leukocytes in the submucosal plexus of the acute DSS paradigm had returned to normal levels following the two-month recovery period (**Supplemental Figure 3A**). Remarkably, however, αSyn accumulation in the ganglia of the submucosal plexus was still almost doubled when compared to αSyn transgenic mice that were not exposed to DSS, and this was exacerbated in αSyn transgenic mice deficient for Cx3cr1 (**Supplemental Figure 3B**). This suggests that accumulation of αSyn is not a transient effect or response and that improper function of monocytes/macrophages contributes to aggravation of this accumulation.

### Experimental colitis-induced at a young age exacerbates αSyn brain pathology and dopaminergic neuron loss in old αSyn transgenic mice

The previously highlighted hypothesis by Braak and colleagues associates αSyn brain pathology in PD with αSyn pathology in the ENS earlier in life ^3,41^. To assess development of brain αSyn pathology and to link it again to IBD risk, we exposed 3-month old hemizygous (Thy1)-h[A30P]αSyn transgenic mice to DSS or normal drinking water and after 23 days on this chronic increasing dose paradigm returned all mice normal drinking water for the rest of their life (Figure 2C). We chose to use the αSyn transgenic model rather than wild type mice for this study because we knew that the model exhibits some αSyn brain pathology that develops slowly also under baseline conditions. After aging up to 9 or 21 months (i.e. mice aged for an additional 6 or 18 months after the chronic DSS paradigm at the age of 3 months, respectively), we analyzed brain regions for pathological αSyn (proteinase K resistant, pSer129-αSyn immunoreactive inclusions). When we examined the αSyn transgenic mice exposed to DSS at 3 months of age, left to live only until to 9 months, we found that they exhibited extremely low levels of pathological αSyn inclusions in the brain, similar to the levels seen in 9 month old hemizygous (Thy1)-h[A30P]αSyn transgenic mice never exposed to DSS (Figure 7 **and Supplemental Figure 4**). Similarly, twenty-one-month old hemizygous (Thy1)-h[A30P]αSyn transgenic mice that only received water during their lifetimes showed relatively low levels of pathological αSyn in the brain (Figure 7 **and Supplemental Figure 4**), which is consistent with previous observations ^42^. In marked contrast, the 21-month-old hemizygous (Thy1)-h[A30P]αSyn transgenic mice that were exposed to DSS at 3 months of age presented with pSer129-positive αSyn pathology throughout various brain regions in a much exacerbated fashion than mice that were aged up to 21 months without having experienced DSS colitis at young age. The significant aggravation of αSyn pathology also in the substantia nigra (p≤0.01 in a negative-binomial mixed-effects model adjusting for multiple comparisons performed over all brain areas) was accompanied by a significant loss of tyrosine hydroxylase (TH) and Nissl positive cells at 21 months of age (p≤0.05, Student’s T-test; Figure 8). Together, we found that DSS colitis at a young age caused an age-dependent exacerbation of αSyn inclusion pathology and a loss of nigral dopaminergic neurons in the brains of αSyn transgenic mice.

**Figure 7.**
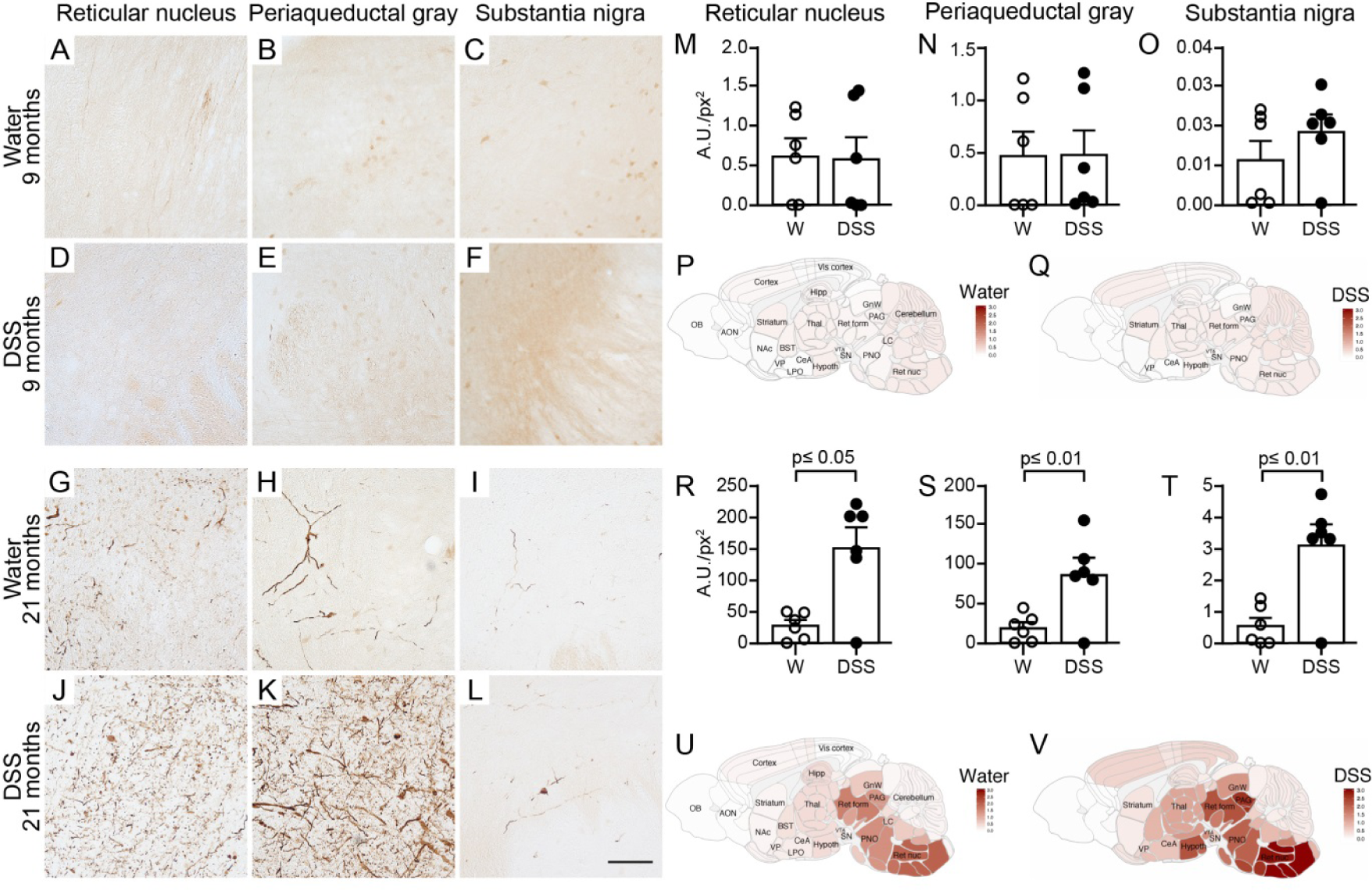
A single chronic DSS colitis insult causes an age-dependent accumulation of proteinase K resistant pSer129-αSyn in various brain regions of (Thy1)-h[A30P]αSyn transgenic mice. A 3-week chronic increasing dose DSS paradigm was performed with 3-month old (Thy1)-h[A30P]αSyn transgenic mice. After recovering and further aging, various brain regions were analyzed for proteinase K resistant pSer129-αSyn immunoreactivity in 9-month **(A-F)** and 21-month old **(G-L)** mice, respectively. The dark brown features in G-L indicate proteinase K resistant pSer129-αSyn. They are barely visible in A-F. Densitometric quantification of pSer129-αSyn immunoreactivity in different brain regions in 9-month **(M-O)** and 21-month old mice **(R-T)** (n=6 mice per group). The two orders of magnitude different y-axes between M-O and R-T confirm the visual impression in panel A-L. Statistical analyses were performed using negative-binomial mixed-effects models adjusting for multiple comparisons. Representative heatmap of the average distribution scores of pSer129-αSyn immunoreactivity for each treatment group in varying brain regions in all the 9-month **(P-Q)** and 21-month old **(U-V)** mice was generated in a sagittal mouse brain (n=10 mice per group). Scale bars: 500 μm.

**Figure 8.**
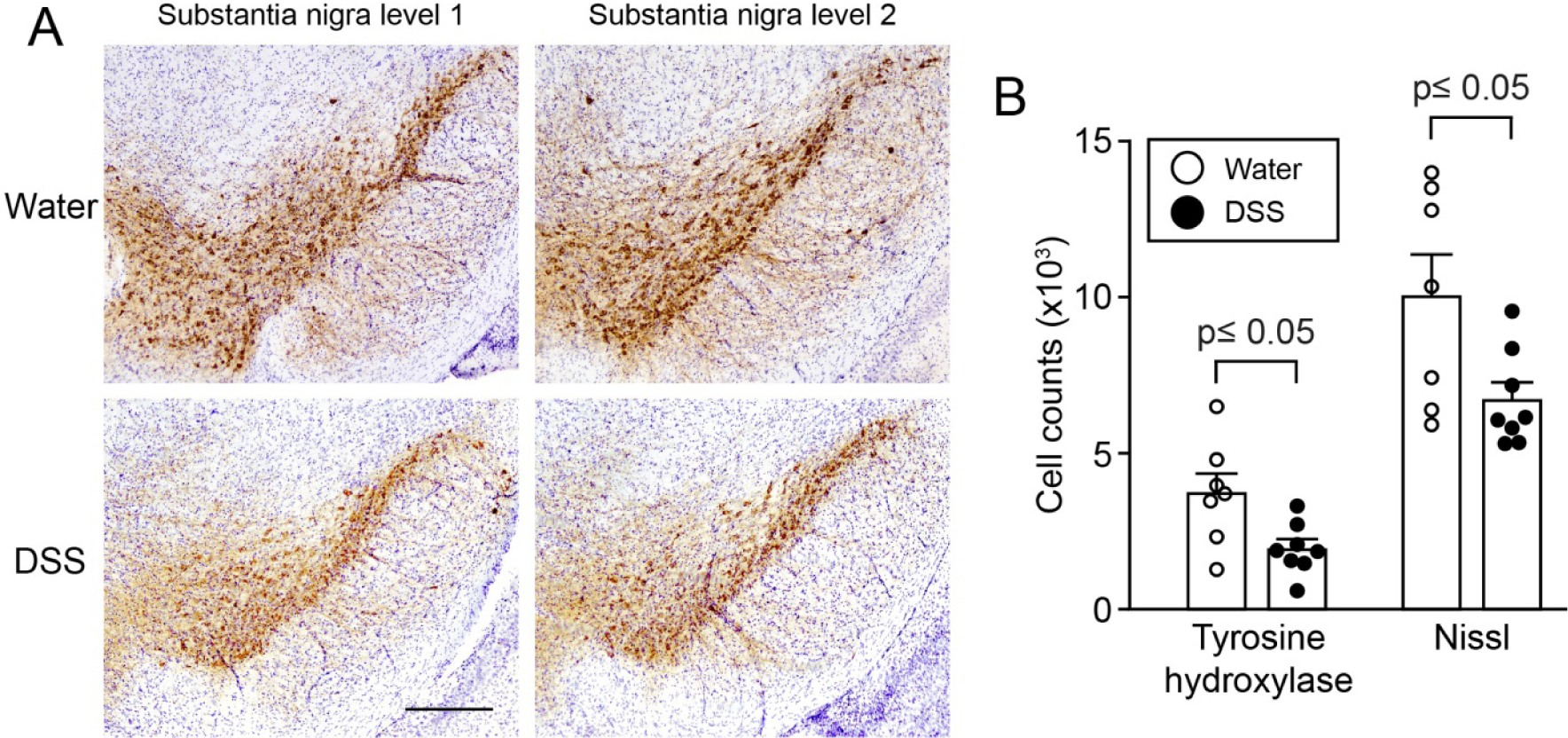
Loss of tyrosine hydroxylase and Nissl positive cells in the substantia nigra of (Thy1)-h[A30P]αSyn transgenic mice at 21 months of age, 18 months post recovery from DSS colitis. (Thy1)-h[A30P]αSyn transgenic mice that were exposed to a chronic DSS-colitis paradigm at 3 months and were aged to 21 months showed a significant loss of mean count of cells with tyrosine hydroxylase (TH) immunoreactivity and cellular Nissl staining in the substantia nigra compared to age-matched littermate mice in the group that did not experience DSS colitis (water). **(A)** Representative images of two levels of the substantia nigra in one mouse per group. **(B)** Stereological quantification of cells positive for TH or Nissl (n=7-8 mice per group). Statistical analyses of the TH dataset were performed using Student’s T-test, while Welch’s T-test was used for the Nissl dataset to adjust for unequal variances. Scale bar: 500 μm.

## Discussion

Currently, there is no therapy for PD available to slow or stop its progression and an obstacle in the quest to develop one is that we do not understand how the disease develops ^43^. Intraneuronal accumulation of αSyn (i.e. Lewy bodies and neurites) is a key neuropathological hallmark and the distribution of Lewy pathology in postmortem brain is used for staging in PD ^2,44^. Accumulation of αSyn has also been observed in the peripheral nervous system in PD, some individuals at risk of developing the disease, and normal individuals ^45–47^. Similar to this finding in people, αSyn-immunoreactive inclusions have also been detected in the ENS of a transgenic mouse model prior to changes in the brain ^48^. Based on preclinical models and postmortem pathology in various organs including the brain, it has also been suggested that αSyn pathology propagates temporospatially in a prion-like manner ^3,44,49–51^. However, the initial factors triggering αSyn aggregation are yet to be established ^43^ and the involvement of peripheral stimuli in the aggregation and pathogenic spread of αSyn is only beginning to unravel.

In this study, we provide evidence that patients with IBD have increased αSyn accumulation in the ENS (Figure 1) and that DSS colitis, i.e. an experimental IBD-like inflammation, triggers αSyn accumulation in the ENS of wildtype mice and in a transgenic mouse model of PD (Figure 3). Interestingly, in IBD patients and in the mouse model of IBD, we observed macrophages filled with αSyn in the inflamed colonic wall (Figure 5). We found aggravation of enteric αSyn accumulation in αSyn transgenic mice lacking Cx3cr1 signaling and amelioration of inflammation and enteric αSyn load by systemic IL-10, suggesting that monocytes/macrophages modulate the process (Figure 4 **and** 6). We further observed that the aggravated αSyn accumulation in the ENS persisted even after two months of recovery from DSS colitis and was aggravated in the absence of CXCR1 signaling, indicating that the effect is not transient and mediated by monocytes/macrophages (**Supplemental Figure 3**). Remarkably, 18 months but not 6 months post induction of DSS colitis (thus, at ages 21 months but not 9 months, respectively), αSyn transgenic mice had developed Parkinson-like brain pathology (Figures 7, 8, **and Supplemental Figure 4**). This included elevated proteinase K resistant pSer129-αSyn pathology in the midbrain, including the substantia nigra, and other brain regions and an average decrease of 30-50% of TH- and Nissl-positive cells in the nigra. We chose to perform the long-term experiments in αSyn transgenic rather than wild type mice. These particular αSyn transgenic mice had previously been shown to slowly develop αSyn pathology in the brain ^33^ making them ideal when asking the question whether transient colonic inflammation can aggravate brain pathology in a genetically predisposed animal. In future long-term studies, we plan to address whether αSyn pathology develops also in the brains of wildtype mice if they sustain transient experimental IBD at a young age. In our experiments presented here, colitis in αSyn transgenic mice recapitulated the accumulation of enteric αSyn which is proposed to occur in humans several years before PD diagnosis ^32^. Additionally, the subsequent age-related development of αSyn pathology together with the loss of nigral dopaminergic neurons in the brain of αSyn transgenic mice mimicked a progression of the disease similar to what is considered to occur in PD.

We established that a mechanism by which peripheral inflammation promotes αSyn accumulation in the colon potentially involves monocytes and macrophages. Both peroral DSS and intraperitoneal LPS administration provoked strong local immune reactions resulting in leukocyte infiltration into the submucosa of the colon. The region of the colon which was inflamed contains the submucosal plexus and is anatomically separated from the myenteric plexus by a thick circular muscle (Figure 2). This discrete localization of inflammation to the submucosa might explain why αSyn only accumulated in the nerves of the submucosal plexus and not in the myenteric plexus of our mice given DSS. The mechanism underlying how intraperitoneally administered LPS leads to submucosal leukocyte infiltration probably involves the monocyte attractant chemokine CCL2 (Figure 4), but the specifics remain to be clarified ^52^. Indeed, CCL2 was upregulated in the colon of our DSS model. However, in contrast to intraperitoneal LPS, where infiltrating macrophages were present in discrete patches in the colonic wall, DSS-related macrophage infiltration was distributed both in small groups and larger randomly distributed patches of cells across the entire colonic submucosa. Also, perorally administered DSS destroys the mucosa of the colon, similar to some forms of ulcerative colitis, resulting in the transient disintegration of the intestinal epithelial barrier. In our (Thy1)-h[A30P]αSyn transgenic mice, the subsequent immune response to the infiltration of commensal bacteria evoked an elevated expression of cytokines such as IL-1β and IL-6. This upregulation was absent in the LPS paradigm in which the intestinal mucosa remained intact. By acting on tight junctions, IL-1β and IL-6 can increase intestinal barrier permeability (gut leakiness), facilitating the recruitment of additional immune cells to the site of the inflammation, eventually culminating in widespread immune activation ^53,54^. Consistent with the breach of barrier permeability in our mouse model, some PD patients exhibit increased colonic cytokines such as IL-1β, IL-6 and TNF, occurring together with increased intestinal permeability ^20,55^. In this context, it is also notable that Crohn’s patients present with increased enteric αSyn expression ^40^ and even more striking that IBD patients on anti-TNF therapy have a reduced risk of developing PD compared to IBD patients not given this treatment ^29^. Here we demonstrate that patients with IBD present with accumulation of αSyn in the ENS, as well as in infiltrating leukocytes nearby. Notably, mucosal macrophages with intralysosomal αSyn content were previously described in the intact human appendix ^56^. These macrophages were in close proximity to the axonal varicosities of the vermiform appendix which showed an enriched staining for αSyn in the mucosal plexus. Furthermore, we recently found that the vermiform appendix contains aggregated and truncated αSyn that has the propensity to seed aggregation of recombinant αSyn *in vitro* ^47^. What could be a functional role of the αSyn species found in abundance in the gut wall? Monomeric and oligomeric αSyn species reportedly act as chemoattractants for neutrophils and monocytes, enhancing the maturation of dendritic cells in the ENS ^19,57^. With such a role in intestinal immunity, it is possible that the tissue destruction induced by DSS in the present study led to release of αSyn, which perhaps served as a chemoattractant for monocytes. The increased abundance of extracellular αSyn and altered intestinal permeability, along with the DSS-evoked inflammatory response may have provided an enabling milieu allowing further αSyn accumulation in the ENS of the colon ^58^. Macrophages and other immune cells are also regulated by several genes including *LRRK2*, an established risk gene for PD and IBD. It will be interesting to explore how mutations in genes that control autophagy, including the *LRRK2* gene, influence the handling of αSyn by macrophages that invade the inflamed colon in our DSS colitis paradigm. Despite the intriguing translational aspect of our finding in the DSS paradigm, others have very recently reported that DSS colitis in mice down-regulates the expression of enteric αSyn on protein levels *in vivo* ^59^. This is in contrast to our immunofluorescence (e.g. increased accumulation of αSyn in submucosal plexus upon DSS colitis; Figures 3, 4, **and** 6) and gene expression data (e.g., no change in endogenous and transgenic αSyn upon DSS colitis; **Supplemental Figure 2**) in the same paradigm and may reflect the well-known lab-to-lab variability that can occur for the DSS models ^60^.

Perhaps the most striking finding in our study was that a single period of DSS-induced colitis at a young age led to an exacerbation of αSyn pathology in the brain of αSyn transgenic mice much later in life (Figure 7). How does the severe αSyn inclusion pathology develop in the brain of these mice? One hypothesis is that the brain αSyn pathology observed in this study could be due to direct effects of peripheral immune activation on the brain and that certain peripheral triggers can directly affect microglial activity. For instance, short-chain fatty acids derived from gut microbiota appear to influence function and maturation of microglia in the mouse brain ^61^ and inflammatory mediators released by gut microbiota into the bloodstream have been suggested to induce brain pathology and behavioral changes in an αSyn transgenic mouse model ^62^. Moreover, rats and nematodes have been reported to develop αSyn inclusions after exposure to the bacterial amyloid protein curli, a protein which stimulates microgliosis, astrogliosis, and secretion of IL-6 and TNF ^63^. Intriguingly, a recent study reported that peripherally applied inflammatory stimuli induce acute immune training (that exacerbates β-amyloid pathology) and immune tolerance in the brain that reprograms microglia, an effect which can persist for at least six months ^64^. Whether this is a relevant mechanism in the DSS paradigm needs to be explored.

Another hypothesis is that the brain αSyn pathology observed may have accumulated following the transfer of pathogenic αSyn seeds from the gut via the vagal nerve. Several experimental studies have demonstrated that pathogenic αSyn seeds can be transferred from the peripheral to the central nervous system. Aggregated recombinant αSyn injected intraperitoneally, intramuscularly or into the gastric wall of certain mouse models of PD results in αSyn inclusions in the brain ^16,65^. Data from animals injected with αSyn protein in the gut wall or viral vectors expressing αSyn into the vagal nerve suggest that pathogenic seeds can be transmitted via the vagal nerve ^15,66^. A role for the vagal nerve in PD was also suggested by an epidemiological study indicating that vagotomy in a Danish population is associated with decreased PD risk ^67^, although this association has been challenged ^68^. In the present study, αSyn pathology was much more prominent in the reticular nucleus (including the vagal area) and midbrain areas (compared to the rostral areas) at 18 months post DSS colitis. Although we did not conduct the definitive experiment of cutting the vagal nerve, our data support the growing body of evidence that the vagal nerve is involved in the accumulation of αSyn aggregates in the brain.

In summary, here we report that individuals with IBD exhibit αSyn accumulation in the colon concomitant with infiltrating monocytes/macrophages positive for αSyn. We also show that αSyn accumulates in the colon of αSyn transgenic and wildtype mice subjected to DSS colitis and that this process is modulated by monocyte/macrophage-related signaling. We further demonstrate that chronic DSS colitis in young αSyn transgenic mice leads to a markedly exacerbated accumulation of αSyn aggregates in the brain when the mice age. In the same aged mice, the numbers of TH- and Nissl positive neurons in the substantia nigra are reduced, suggestive of a neurodegenerative process.

Together, our findings are in consonance with studies demonstrating a link between IBD and PD ^28,29,69^ and suggest a critical role for intestinal inflammation and αSyn accumulation in the initiation and progression of PD.

## Methods

### Mice

Male C57BL/6 wild type mice (Jackson Laboratories, Bar Harbor, USA), hemizygous Tg(Thy1-SNCA*A30P)18Pjk ((Thy1)-h[A30P]αSyn) ^33^ and Tg(Thy1-SNCA*A30P)18Pjk crossed with Cx3cr1tm1Litt ((Thy1)-h[A30P]αSyn /CX3CR1-def; homozygous for Cx3cr1-GFP knock-in allele; ^70^ transgenic mice were used for the study. (Thy1)-h[A30P]αSyn transgenic mice express mutant human αSyn under the neuron selective Thy1 promoter. (Thy1)-h[A30P]αSyn transgenic mice were crossed to Cx3cr1-def transgenic mice which express eGFP replacing fractalkine gene expression. All mice were maintained on a C57BL/6 background for more than 10 generations and under specific pathogen-free conditions. To the extent possible, littermates were used in the experiments. Health status was monitored daily during experiments.

### Human Subjects

Samples from patients with Crohn’s disease (CD), ulcerative colitis (UC) or healthy subjects (HS) were provided by the tissue bank, Institute of Pathology, University of Bern. Briefly, specimens were obtained from patients who underwent surgical procedures at the University Hospital (Inselspital) in Bern, Switzerland between 2004 and 2011. Three selected male patients previously clinically diagnosed with UC with a reported disease duration > 6 year (n=3) and undergoing steroid therapy combined with either metronidazole or mesalazine. CD patients were of mixed gender and aged 22-56 years ranging from 2 months to 11 years post disease diagnosis undergoing treatment with either infliximab or mesalazine in combination with steroids. Healthy subjects were of mixed gender with no report of inflammatory bowel disease, aged 40-59. All samples contained the mucosa and submucosa regions including minor parts of the circular muscle layer. Following surgical removal, tissue samples were immediately immersed in O.C.T. compound (VWR International GmbH, Dietikon, Switzerland), frozen in liquid nitrogen and stored at −80°C. Diagnosis of disease status was made according to established criteria for histopathological analysis.

### Experimental IBD in mice with DSS and LPS

Paradigms for the induction of inflammation were either 1 week (acute) or 3-4 weeks (chronic) with or without an incubation phase of 2, 6 or 18 months post application (Figure 2). Acute systemic inflammation was induced by intraperitoneal Lipopolysaccharide (LPS) application ^71^ of 0.5 mg/kg in 100 µl injection volume on day 1 and 4 (Sigma-Aldrich Chemie GmbH, Steinheim, Germany, LPS 055:B5). Acute colitis was induced by application of 36-50kDa Dextran Sulfate Sodium (DSS) ^72^ (160110, MP Biomedicals, LLC, Illkirch, France) at 0%, 1%, 2.5% or 5% in autoclaved drinking water for 5 continuous days respectively, followed by 2 days of water (1 DSS application cycle). Chronic colitis was achieved by 4 repeating DSS application cycles. The DSS concentration during 4 weeks of chronic colitis was either 1% or 2.5% for 4 weeks or 2.5%-4% raised 0.5% every week for 4 weeks. Mice from same littermate group were randomized per cage into vehicle and inflammation inducing agent.

### IL-10 treatment and exposure measurement

Two different forms of mouse IgG bound murine IL-10 (mIgG(v1)-mIL10 and mIgG(v2)-mIL10) were diluted in pre-prepared sterile formulation buffer comprised of 0.5% mouse serum supplemented with 25mM citrate, 300mM arginine to a final concentration of 0.75 mg/ml and the pH adjusted to 6.7 on the day of application. Each mouse was treated once with 150 µg i.p concurrently with the initiation of the acute colitis paradigm with 5% DSS. The concentrations of mIgG-mIL10 fusion proteins in murine serum samples were determined by enzyme-linked immunosorbent assays (ELISA) specific for the Fab moiety of the administered mIgG-mIL10 fusion protein. Biotinylated mIgG-mIL10-specific target molecules were used for capturing, goat anti-mIg IgG-HRP conjugate and peroxidase substrate ABTS was used for quantitative detection of mIgG-mIL10 fusion proteins.

### Immunohistochemistry

Animals were injected with a lethal dose of pentobarbital (150 mg/kg). Upon full anesthesia, mice received transcardial perfusion with room temperature phosphate buffered saline (PBS). For biochemical and immunohistochemical analysis, one section of either the proximal colon was fresh frozen and stored at −80°C or post-fixed in 4% paraformaldehyde (PFA) solution for 24 h. Following post-fixation, organs were incubated in 30% sucrose/PBS at 4°C for at least 48 h before further processing. Subsequently, enteric tissue was cryotome-sectioned to 35 µm thick longitudinal sections (approx. 1 cm length). The brain was collected and post-fixed for 24 h in 4% PFA followed by 30% sucrose in phosphate buffer until cryo-sectioning of floating sections at 40 μm. Histological analysis of the colon was performed using standard hematoxylin staining. Immunohistochemical staining was accomplished using the Vectastain Elite ABC Kits and Peroxidase Substrate Kit SK-4100 (Vector Laboratories, Burlingame, CA, USA) or fluorescently labelled secondary antibodies (Alexa coupled to dye 488, 555 or 647, Life Technologies, Zug, Switzerland). The following primary antibodies have been used for overnight incubation at a dilution of 1:1000: monoclonal antibody to human α-synuclein (clone 211, sc-12767, Santa Cruz Biotechnology, Heidelberg, Germany; used on tissue from human αSyn transgenic mice), monoclonal antibody to rat α-synuclein but cross-reactive with murine and human αSyn (Syn1/clone 42, BD Transduction Laboratories, Allschwil, Switzerland; used for wild type mice and in human colon), polyclonal antibody to the peripheral neuronal marker Peripherin (Millipore Corporation, Billerica, MA, USA), and polyclonal antibody to macrophage marker Iba1 (Wako Chemical GmbH, Neuss, Germany). To detect phosphorylated αSyn (pSer129 pathology) in the free-floating brain sections, monoclonal antibody (ab51253, Abcam, Cambridge, USA) to human αSyn was used at a dilution of 1:10000. Prior to the pSer129 staining, the free-floating brain sections were incubated for 10 min at room temperature in a phosphate buffered saline solution containing 10 μg/mL proteinase K (Cat # 25530015; Invitrogen, California, USA). TH-immunoreactive cells were detected using a polyclonal antibody (657012, Millipore Sigma) at a dilution of 1:1000. To measure the density of Nissl-positive cells, the TH-stained cells were counter-stained with Cresyl violet. The slides were incubated in 0.1% Cresyl violet solution for 9 min and then dehydrated in 95% and 100% ethanol and then xylene prior to coverslipping with Cytoseal 60 mounting media (Thermo Fisher Scientific). Quantifications of the blind-coded TH/Nissl stained slides were done using Stereoinvestigator (version 2017.01.1; MBF Bioscience, Williams, VT, USA) on Imager M2 microscope (ZEISS) coupled to a computer. We analyzed 5-7 nigral sections per animal, and a total of 7-8 animals per treatment group. We outlined the substantia nigra pars compacta and counted every TH-immunoreactive and Nissl-positive cell in that area and computed the number of cells per section, generating the mean cell density per animal. We then calculated the mean density of cells per treatment group and analyzed the data using unpaired Student’s T-test after confirming normality and homoscedasticity in Prism 7.0 (GraphPad Software).

### Imaging and stereological quantification of αSyn deposits in enteric nervous system

Imaging and stereological quantification was performed on a Zeiss Axio Imager Z2 fluorescence microscope (Carl Zeiss AG, Jena, Germany). Leica TCS SP5 confocal system using an HCX PL APO CS 40x 1.3 oil UV or an HCX PL APO LB 63x 1.4 oil UV objective was utilized for image recording. Accumulation of αSyn in the ENS was assessed on a random set of 3 adjacent 35 µm thick, αSyn-immunostained sections comprising the myenteric and submucosal neuronal plexuses. Analysis was performed with the aid of Stereologer software (Stereo Investigator 10, MBF Bioscience, Williams, VT, USA) as described previously ^73^. In the myenteric plexus ganglion volume was defined by multiple outlined plexuses containing a range of 5-20 neuronal cells and quantified by the optical fraction fractionator technique. In contrast to the myenteric plexus, the submucosa consists of compact plexuses with 1-5 cells including interconnecting neurites. Therefore, the entire submucosa was set as region of interest, analyzed with the area fraction fractionator technique. Results of the submucosal plexus are displayed by percent area containing αSyn deposits. For the IL-10 experiment, αSyn positive inclusions from immunofluorescence images were counted for each image. Inclusion body-like features were filtered based on having a size between 12 and 50000 pixels and a minimal intensity value greater than 300. The filtering step was included to exclude small background features and macrophages (very large spots). The counts were then aggregated to the animal level by summing the inclusion feature counts of all images per animal and then normalizing for (i.e. dividing by) the number of images for a given animal. Upon exploratory data analysis two animals were excluded: one mouse because it only had one image and another due it being an outlier, based on its infiltration score and image data.

### Blinding of experimenters

For analyses of colon and brain tissue on slides, a second individual assigned unique codes to stained slides. Therefore, the experimenter conducted the analyses blinded to the identity of the mice. For randomization of treatment groups see above.

### Quantification of leukocytes infiltration

To determine the leukocyte covered area in the colon after LPS or DSS application, three adjacent hematoxylin stained sections were quantified. Total area of colon sections and localizations of leukocyte assemblies within the tissue architecture were identified and outlined utilizing Stereologer Software (Stereo Investigator 6, MBF Bioscience, Williams, VT, USA). Percentage of leukocyte covered area has been set in proportion to total area of the analyzed colon section. For the IL-10 experiment, hematoxilin stained colon slices were examined by an expert pathologist blinded to treatment conditions. A score of 0-3 was assigned to each section for each of the 3 layers lamina propria, submucosa and muscularis based on the degree of inflammatory infiltration. A score of 0 denoted no inflammation and a score of 3 indicated extensive infiltration. The mean of the values for all 3 layers was taken as the final measure of leukocyte infiltration per mouse.

### Quantification of αSyn/Iba1 double positive macrophages

The number of α-syn+/Iba1+ positive cells was evaluated by quantification of 10 random regions in 2 adjacent sections of the proximal colon. The region of interest was set to contain the myenteric plexus/circular muscle layer and the submucosal plexus. Cells were assessed for positive αSyn staining and concomitant co-localization with the macrophage marker Iba1 was quantified.

### Scoring of pSer129 pathology and brain heatmap

We evaluated pSer129 pathology on a full series of immunostained coronal sections from 10 mice per treatment group (i.e. water vs. DSS-treated groups) on blind-coded slides using a previously described method ^74^. We visualized pathology from one hemisphere of all brain sections (apart from the olfactory area) using NIKON Eclipse Ni-U microscope and assigned scores ranging from 0 to 4 to each brain area based on the relative abundance of PK-resistant pSer129-positive inclusions (i.e. cell bodies and neurites). In this case, 0 = no aggregates, 1 = sparse, 2 = mild, 3= dense, 4 = very dense. For the heatmap, we obtained the average score values of each brain area for each treatment group. The average data for each treatment group (n=10/ group) was then represented as a heatmap in a sagittal mouse brain background (http://atlas.brain-map.org/atlas?atlas=2#atlas=2&structure=771&resolution=16.75&x=7755.7470703125&y=3899.625&zoom=-3&plate=100883867&z=5).

### Densitometry of pSer129 αSyn brain pathology

The density of pSer129 pathology in 12 major brain areas (reticular nucleus, pontine reticular nucleus, periaqueductal gray, gray and white layer, reticular formation, substantia nigra, ventral tegmental area, thalamus, hypothalamus, central amygdala, pallidum and striatum) was determined in the water and DSS-treated animals. A NIKON Eclipse Ni-U microscope was used to acquire 20x magnification images (without condenser lens) from all the indicated brain areas, using the same exposure time for all images. In all cases, images were acquired on three sections separated by 420 μm intervals (localized between Bregma). We then processed the acquired images using Image J64 ^75^, created a mask (to exclude background) that redirects to the original image for analysis, measured the total area and the mean grey value of the area that had inclusions. For brain areas such as periaqueductal gray that do not fill the entirety of the field to be analyzed, we drew a contour of the area and the analysis was performed only within that contoured area. We subsequently calculated the grey value of the area per square pixels for each image (i.e. A.U./px^2^ = mean grey value x area stained/total area assessed). Based on this, we calculated the average grey value per square pixels for each brain area for each animal (*n* = 6 mice/group), and then extended this calculation to determine the average grey value per square pixels for each treatment group and each of the twelve brain areas of interest.

### mRNA expression

To assess mRNA expression levels from the proximal colon, RNA was extracted from fresh frozen tissue with MagnaLyser green beads (Roche Diagnostics, Mannheim, Germany) and Qiazol Lysis (Reagent cat.no.79306, Hilden, Germany) purified on MagnaPure LC (HP Kit no.03542394001, F. Hoffmann - La Roche AG, Rotkreuz, Switzerland) and amplified via real-time PCR (4ng RNA/reaction; Lightcycler 480, Roche Diagnostics Corporation, Indianapolis, USA). Amplification of mRNA was performed by using TaqMan probes for human or murine specific α-synuclein and for selected cytokines/chemokines (Applied Biosystems Europe B.V., Zug, Switzerland). Target mRNA was normalized to tissue-specific murine GAPDH levels and displayed as relative expression after 30 amplification cycles.

### Statistics

Statistical analysis of gut pathology and inflammation was performed using GraphPad Prism 6.04 or 7.0 software (GraphPad Software, Inc. La Jolla, CA, USA). The results are expressed as mean values ± standard errors of the mean (SEM). Student’s T-test (or Welch’s T-test for unequal variances) was used to compare two groups and ANOVA was used for multi-comparison of groups followed by Tukey HSD post-hoc analysis. For the statistical analysis of the pSer129 αSyn brain pathology, negative-binomial mixed-effects models with a random intercept for each sample were used to analyze the dataset via the ‘lme4’ (http://lme4.r-forge.r-project.org/) package in R v 3.4.4. To analyze the pSer129 αSyn cell count dataset, an offset for the total area examined was included to model the densities. Linear contrasts with false discovery rate (FDR) adjustments were then used to test our hypotheses and account for multiple testing (for brain area and experimental group). Like the pSer129 dataset, the Iba-1/αSyn-double positive dataset were analyzed using negative-binomial regression and Tukey HSD adjusted contrasts to test our hypotheses.

For the statistical analysis of the mRNA expression, data quality was assessed by inspecting the distribution of Cp values of reference endogenous genes across samples, by inspecting the level of Cp variation between technical replicates and by exploring the samples multivariate signal distribution as in a principal component analysis. Relative gene expression levels were expressed as 2^-(Cpgene–CpRef)^. Statistical analyses to assess the effect of the experimental conditions on the log2 gene expression levels were done with linear models using the *limma* package (Bioconductor/R, Smyth, 2005). These analyses were implemented in R v3.1.1.

For the statistical modelling of the effects of the IL-10 treatment on αSyn counts, as well as infiltration scores, the levels for IgG1(v1)-IL10 and IgG1(v2)-IL10 treatment were compared to the positive (vehicle/DSS) control. Additionally, since levels of the control antibody treatment (IgG1(v1)) were very similar to the positive control, the two groups were pooled in further contrasts in which effects of individual antibodies or control IgG was assessed. For αSyn counts, a linear model on the treatment groups with one-degree freedom contrasts was applied. For the infiltration score a Kruskal-Wallis test, with the same contrasts, was used.

### Study approvals

The human subjects’ study was conducted with the approval of the local Ethical Committee in Bern No. 47/04. Written informed consent was obtained from each patient. The animal experiments were approved by a Roche internal review board and the local authorities.

## Author contributions

S.G., N.M. and L.S. planned and performed the in vivo experiments, colon immunostaining, analysis, and quantification; S.G. and N.M. drafted a first version of the manuscript; E.Q. performed, imaged, quantitated pSer129, TH and Nissl staining in the brain sections, and drafted a more advanced version of the manuscript with J.A.S., who also provided helpful discussion. F.B. and K.O.S. supported the image acquisition and image analysis for the colon samples; M.St. performed imaging and data analysis of experiments with wildtype mice; G.D.P. and J.S.P. performed statistical analysis of the DSS experiments; H.R. and M.H. performed mRNA analyses; M.Se. trained S.G. and L.S. on mouse necropsy and supported their work; P.M. performed expert pathology staging on leukocyte infiltration; T.E. and A.W. provided mIgG-mIL-10 fusion proteins and measured serum exposure; Z.M. performed statistical analysis for the pSer129 αSyn immunohistochemistry data. M.L.E.G. provided helpful discussion and project planning. A.H. co-mentored S.G. and N.M., performed expert pathology staging on leukocyte infiltration and contributed to experimental planning. C.M. trained S.G. on the colitis model, provided human tissue and expert input on the experimental IBD model. M.B. and P.B. co-mentored Roche Postdoctoral Fellows S.G. and N.M., conceived and oversaw the study, and performed experimental planning; M.B., P.B. and E.Q. wrote the final version of the manuscript.

## Acknowledgments

We acknowledge the human donors for providing tissue used in this study. We thank Drs. L. Ozmen, A. Bergadano, and A. Su for their tremendous support in maintaining the mouse colony and establishing of relevant animal experiment licenses, and we are grateful to the animal care takers, veterinarians and many unnamed staff at Roche for their valuable work with the mice in this study. In addition, at Roche we thank Dr. K.G. Lassen for critical input to the paper, Dr. C. Ullmer for co-mentoring S.G. and providing scientific input, Dr. L. Collin for helping with confocal imaging and we are grateful to Dr. T. Kremer, N. Haenggi, D. Mona, A. Girardeau, and J. Messer for providing support in tissue dissections and G. Walker and R. Lauria for technical support. Ms. E. Schulz from VARI assisted with immunostaining of the brain tissue. We thank the Contract Research Organization Frimorfo for carefully sectioning the brains for this study. We acknowledge Drs. L. Gaudimier (née Chicha) and F. Pan-Montojo for scientific discussions early in the project and Dr. W. Zago from Prothena for valuable scientific input throughout the project. P.B. reports relevant grants from NIH (R01DC016519-01, 1R21NS106078-01A1 and 5R21NS093993-02), Department of Defense (W81XWH-17-1-0534), The Michael J. Fox Foundation for Parkinson’s Research, and Cure Parkinson’s Trust. Finally, we thank the Roche Postdoctoral Fellowship Program for supporting S.G. and N.M.

## Conflicts of Interest

At the time of the study S.G. and N.M. were Roche Postdoctoral Fellows employed by Roche and L.S., F.B., G.D.P., J.S.P., K.O.S., H.R., M.H., M.Se. M.St., P.M., A.W., T.E., A.H. and M.B. are or were fulltime employees or trainees at Roche and they may additionally hold Roche stock/stock options. S.G. and L.S. are currently employees of Neurimmune AG, Schlieren, Switzerland. P.B. has received commercial support as a consultant from Renovo Neural, Roche, Teva, Lundbeck A/S, AbbVie, NeuroDerm, Fujifilm Cellular Dynamics, Living Cell Technologies, IOS Press Partners, and Axial Biotherapeutics. Additionally, P.B. has received commercial support as a consultant from Renovo Neural, Roche, Teva, Lundbeck A/S, AbbVie, NeuroDerm, Fujifilm Cellular Dynamics, Living Cell Technologies, IOS Press Partners, Axial Biotherapeutics and CuraSen. P.B. has received commercial support for grants/research from Renovo, Roche, Teva, and Lundbeck and has ownership interests in AcouSort AB. The other authors do not have conflicts of interest with regard to this research.

## Notes

#### Summary of Updates

Edited and updated entire manuscript (text, new references, figures) for submission to another journal. Content remains the same.

