## Supplemental Figures for "Experimental colitis drives enteric alpha-synuclein accumulation and Parkinson-like brain pathology"

Grathwohl et al., Figure S1

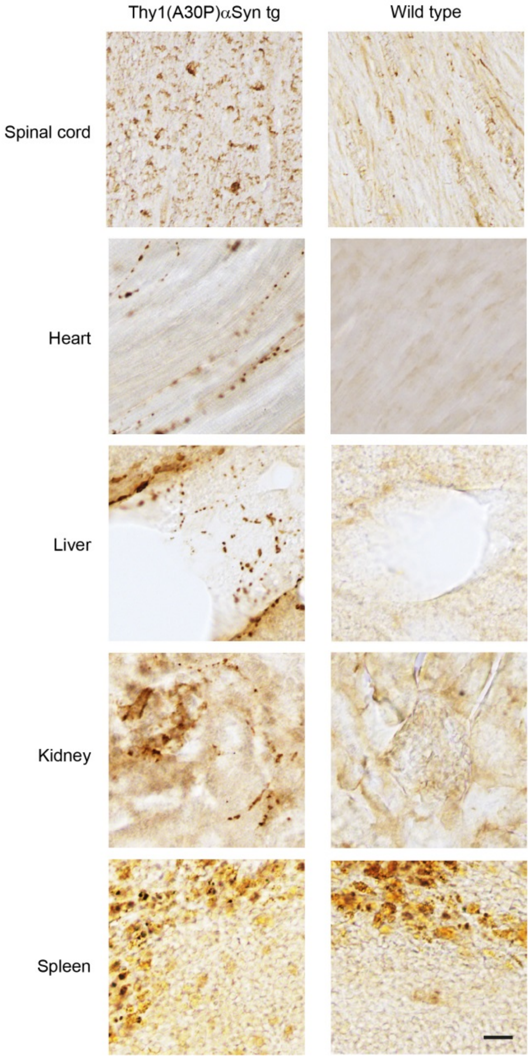


**Supplemental Figure 1**

**Alpha-synuclein is expressed in majority of organs in wildtype and (Thy1)-h[A30P]αSyn transgenic mice.** Immunohistochemical detection of human αSyn (clone LB509 monoclonal antibody) in (Thy1)-h[A30P]αSyn transgenic mice or of endogenous murine αSyn in wild type mice (syn1 monoclonal antibody) in various organs. Note the typical dot-like structures of the human αSyn in the transgenic mice reminiscent of neuritic inclusions and the very low expression of endogenous murine αSyn in the wildtype mice. The pronounced brownish staining in the spleen is due to the abundant iron which is exposed by the chromogenic staining method. Note, thymocytes in the spleen do not stain for human αSyn supporting the selectivity of the expression of the transgenic human αSyn under the modified Thy1.2 cassette; e.g. not expressed in thymocytes (39). Scale bar 20 µm.

Grathwohl et al., Figure S2

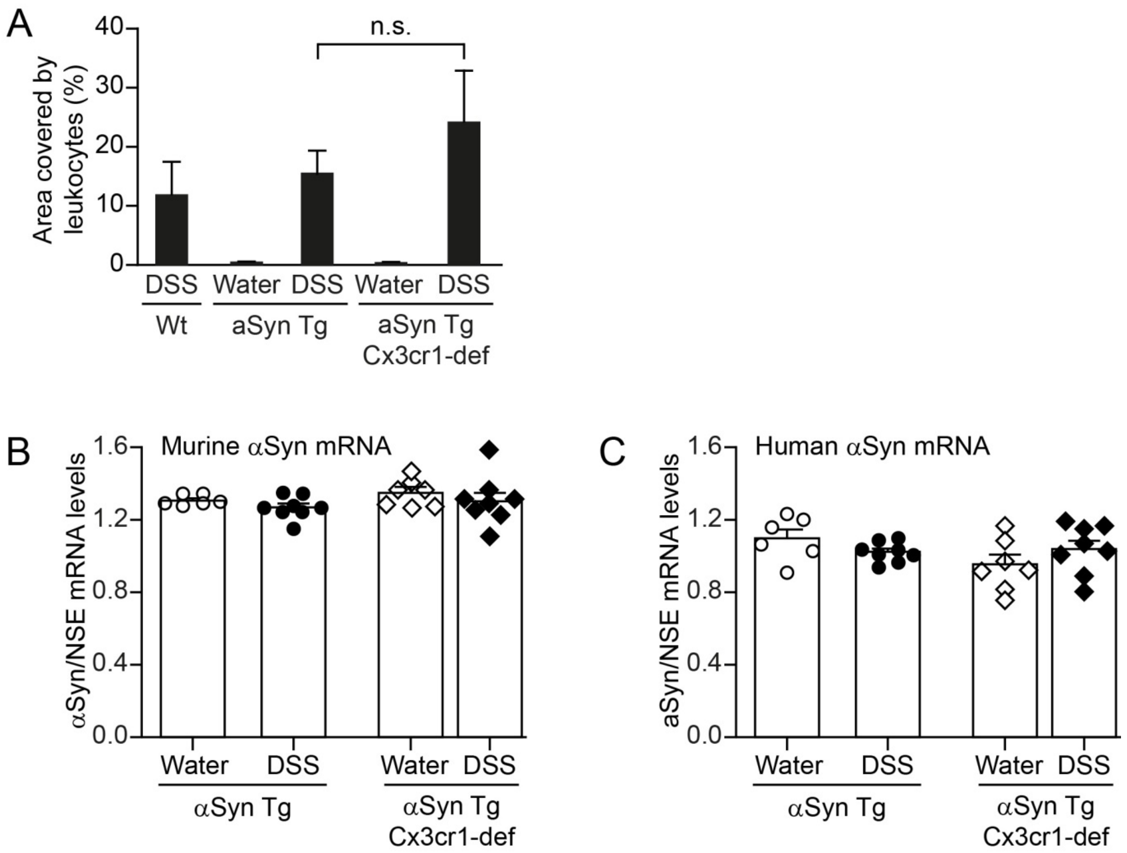


**Supplemental Figure 2**

**Expression of endogenous and transgenic αSyn in the colon is unchanged after acute DSS colitis.** **(A)** Administration of DSS (acute 5%) induces leukocyte infiltration in wildtype, (Thy1)-h[A30P]αSyn transgenic (αSyn Tg) and Cx3cr1-deficient (Thy1)-h[A30P]αSyn transgenic mice (αSyn Tg Cx3cr1-def) (Two-way ANOVA with Tukey post hoc test). Expression levels of endogenous murine **(B)** or transgenic human αSyn **(C)** mRNA were normalized to mRNA levels of the neuronal marker neuron specific enolase (NSE) to correct for potential neuronal loss.

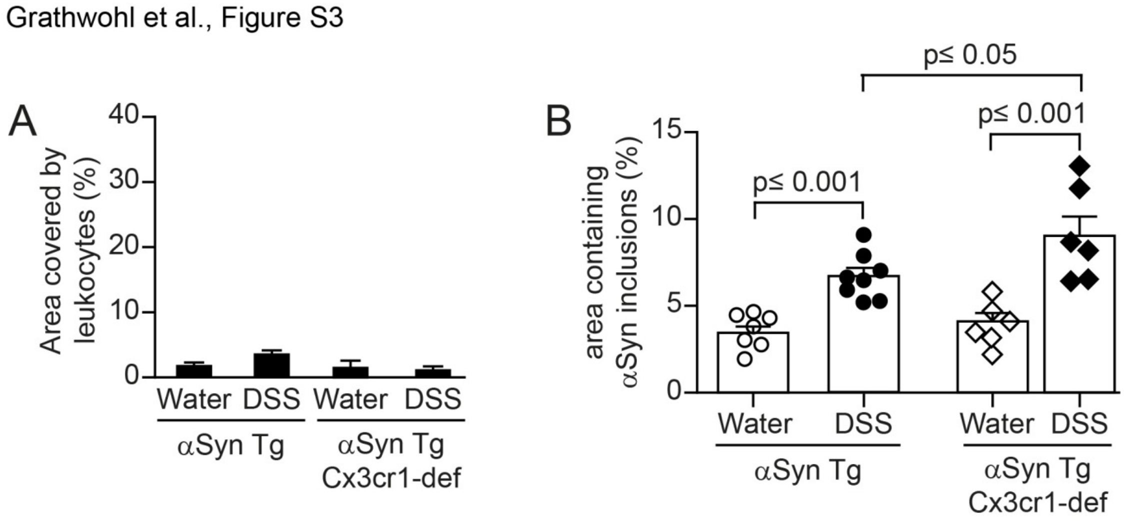


**Supplemental Figure 3**

**DSS colitis induced accumulation of αSyn in submucosal plexus of (Thy1)-h[A30P]αSyn transgenic mice remain stable long after recovery.**

A 4-week chronic DSS paradigm was performed with (Thy1)-h[A30P]αSyn (αSyn Tg) and (Thy1)-h[A30P]αSyn crossed with Cx3cr1-def mice (αSyn Tg Cx3cr1-def). After recovery for 2 months and thus analysis at the age of 6 months the colon was inspected for signs of inflammation (area covered by leukocytes) and amount of αSyn inclusions (area containing αSyn inclusions). Statistical analyses were performed using two-way ANOVA with Tukey post hoc testing.

Grathwohl et al., Figure S4a

Aged up to 9 months (6 months post a 3-week chronic increasing dose DSS colitis paradigm at the age of 3 months)

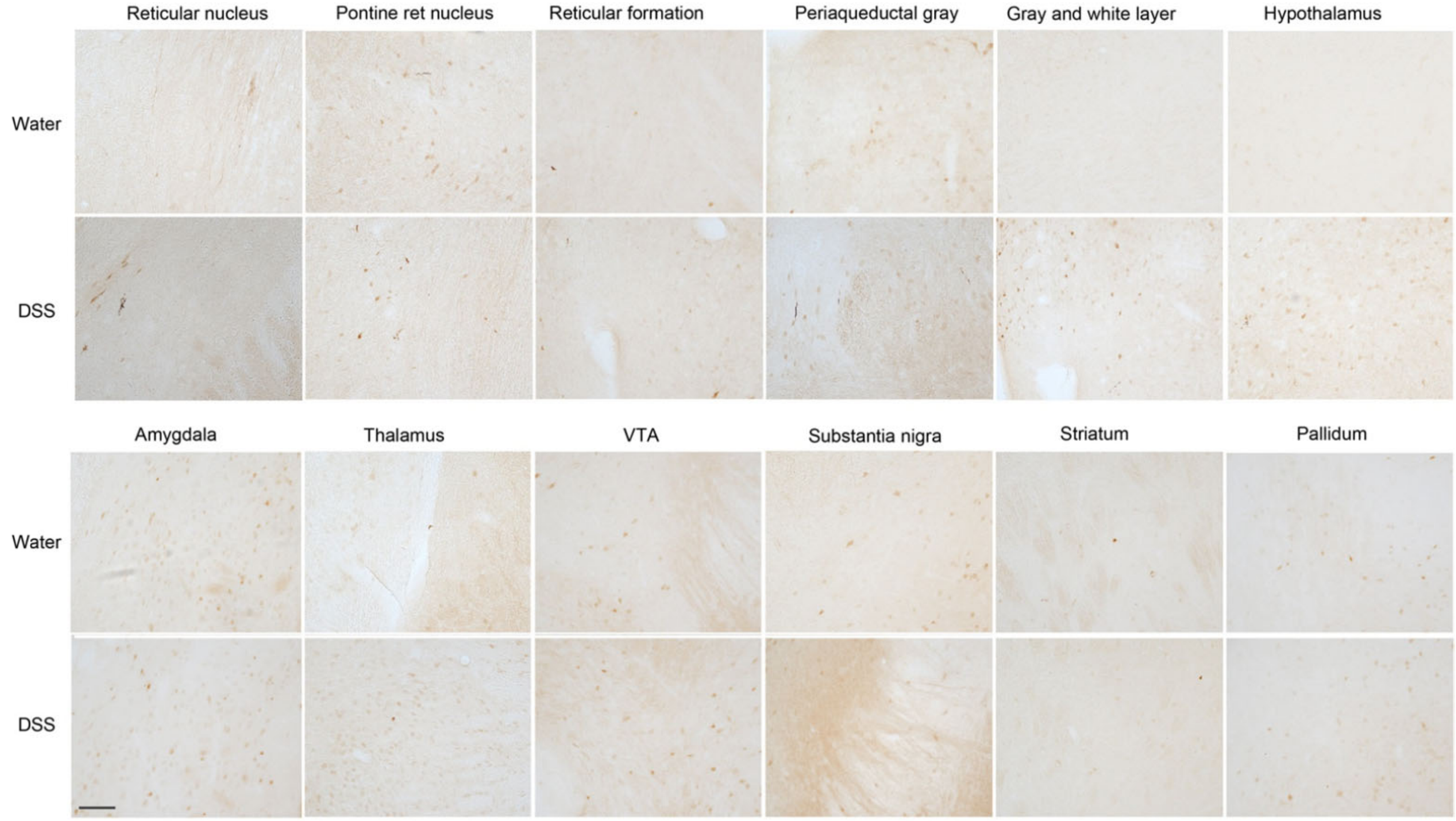

Grathwohl et al., Figure S4b

Aged up to 21 months (18 months post a 3-week chronic increasing dose DSS colitis paradigm at the age of 3 months)

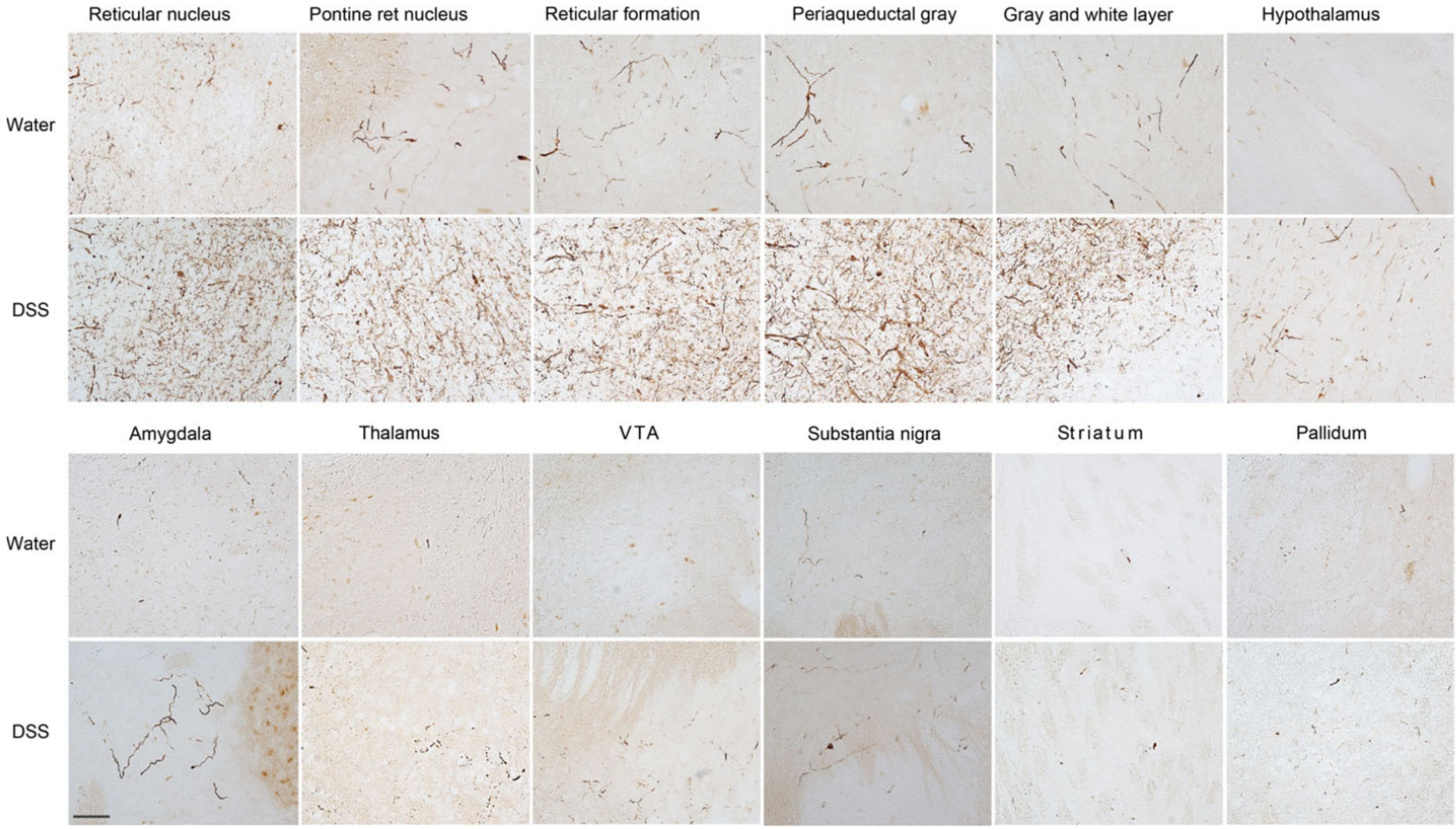

Grathwohl et al., Figure S4c

Quantification of Figure S4b

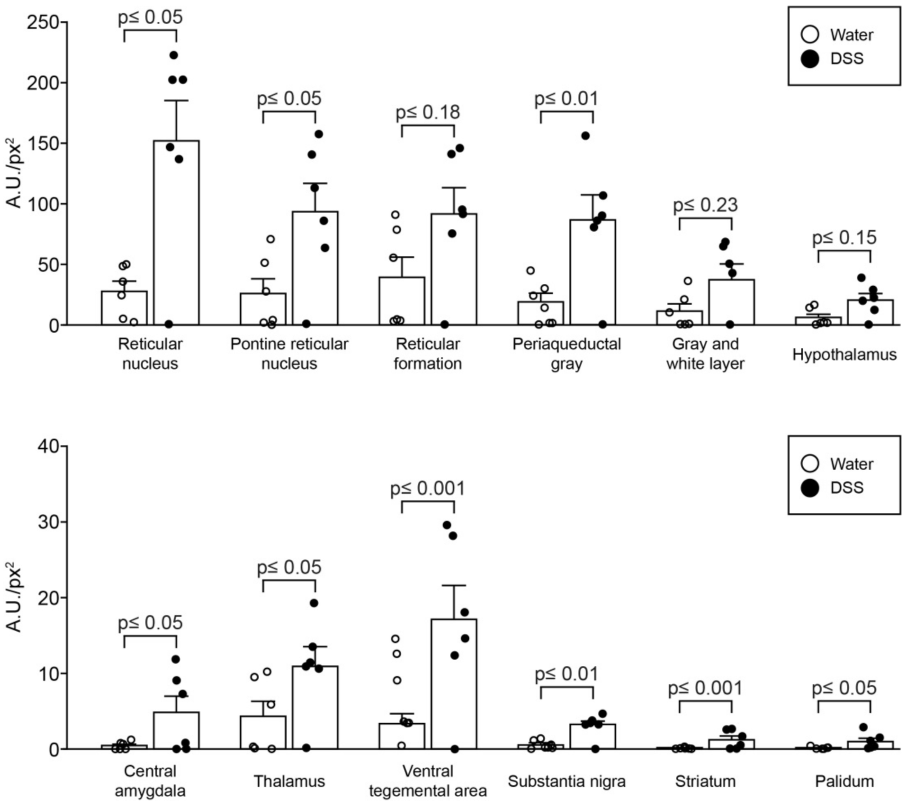


**Supplemental Figure 4(a-c)**

A 3-week increasing dose chronic DSS paradigm was performed with 3-month old (Thy1)-h[A30P]αSyn transgenic mice. After recovery and further aging, various brain regions were analyzed for proteinase K resistant pSer129-αSyn immunoreactivity in 9-month **(a)** and 21-month old **(b)** mice. Densitometric quantification of pSer129-αSyn immunoreactivity in different brain regions in the 21-month old **(S4b)** mice were measured (n=6 mice/group). Statistical analyses were performed using linear mixed-effects model adjusting for multiple comparisons. A.U./px$^2$, = mean grey value x area stained/total area assessed. Scale bars: 500 μm.
